# Simultaneous Membrane and RNA Binding by Tick-Borne Encephalitis Virus Capsid Protein

**DOI:** 10.1101/2022.10.06.511088

**Authors:** L. I. A. Pulkkinen, S. V. Barrass, M. Lindgren, H. Pace, A. K. Överby, M. Anastasina, M. Bally, R. Lundmark, S. J. Butcher

## Abstract

Tick-borne encephalitis virus is an enveloped, pathogenic, RNA virus in the family *Flaviviridae*, genus *Flavivirus*. Viral particles are formed when the nucleocapsid, consisting of an RNA genome and multiple copies of the capsid protein, buds through the endoplasmic reticulum membrane and acquires the viral envelope and the associated proteins. The coordination of the nucleocapsid components to the sites of assembly and budding are poorly understood. Here, we investigate nucleocapsid assembly by characterizing the interactions of the wild-type and truncated capsid proteins with membranes by using biophysical methods and model membrane systems. We show that capsid protein initially binds membranes via electrostatic interactions with negatively-charged lipids which is followed by membrane insertion. Additionally, we show that membrane-bound capsid protein can recruit viral genomic RNA. We confirm the biological relevance of the biophysical findings by using mass spectrometry to show that purified virions contain negatively-charged lipids. Our results suggest that nucleocapsid assembly is coordinated by negatively-charged membrane patches on the endoplasmic reticulum and that the capsid protein mediates direct contacts between the nucleocapsid and the membrane.

## Introduction

Tick-borne encephalitis virus (TBEV) is a viral pathogen in the family *Flaviviridae*, genus *Flavivirus* (Lindenbach *et al*, 2013; Simmonds *et al*, 2017). TBEV is mainly spread by ticks, and causes life-threatening neurological symptoms with human fatality rates as high as 40 % depending on the virus subtype (Bogovic & Strle, 2015; Ruzek *et al*, 2019). Although an effective vaccine exists, the frequency of TBEV infections has increased dramatically in recent decades (Bogovic & Strle, 2015; Ruzek *et al*, 2019). Furthermore, there are no antivirals available, and the treatment of patients is limited to symptomatic care (Bogovic & Strle, 2015; Ruzek *et al*, 2019).

The TBEV virion has a typical flavivirus structure (Füzik *et al*, 2018; Pulkkinen *et al*, 2022). The surface of the virion is made up of two protein species, E and M, that are embedded in the virion envelope in an icosahedrally-symmetric lattice (Füzik *et al*, 2018; Pulkkinen *et al*, 2022). The nucleocapsid (NC) consists of a positive-strand RNA genome of about 11 kb and multiple copies of the capsid (C) protein. However, the structure of the NC inside the particle and its assembly remain poorly understood (Füzik *et al*, 2018; Pulkkinen *et al*, 2018, 2022). The NC assembles at the endoplasmic reticulum (ER) and buds into the ER lumen acquiring both the lipid envelope and the envelope proteins during this process (Lindenbach *et al*, 2013; Pulkkinen *et al*, 2018). This yields immature, non-infectious particles that need to undergo both pH-mediated conformational changes as well as proteolytic cleavage to mature (Stadler *et al*, 1997). In addition to the NC itself, assembly and budding involve at least E, the M protein precursor (prM), the viral non-structural (NS) proteins NS2A, NS2B-NS3 and multiple host factors such as SPCS1, nucleolin, DDX56, and components of the endosomal sorting complex required for transport (ESCRT) (Corver *et al*, 2000; Ferlenghi *et al*, 2001; Lorenz *et al*, 2003; Kümmerer & Rice, 2002; Liu *et al*, 2003; Leung *et al*, 2008; Patkar & Kuhn, 2008; Xu *et al*, 2011; Balinsky *et al*, 2013; Xie *et al*, 2013; Voßmann *et al*, 2015; Xie *et al*, 2015; Blazevic *et al*, 2016; Tabata *et al*, 2016; Reid & Hobman, 2017; Ma *et al*, 2018). Currently, it is not clear if the NC assembles before or concurrently with membrane budding, but since isolated NCs have not been observed in cells, the processes are likely interlinked (Offerdahl *et al*, 2012; Miorin *et al*, 2013; Hirano *et al*, 2014; Yu *et al*, 2014). The central role of the C protein is to recruit genomic RNA to the ER budding site either by interacting with the transmembrane domains of E and prM, host proteins, viral NS proteins, or specific ER lipids (Pulkkinen *et al*, 2018). However, the regulation of these interactions is not understood.

The C protein is translated as part of the single, ER-embedded TBEV polyprotein. It initially has a C-terminal membrane-spanning anchor of 20 residues, which is cleaved by the viral NS2B-NS3 protease (Amberg *et al*, 1994). The mature form of TBEV C protein consists of 96 residues. Homology modelling predicts a similar fold to that of other flavivirus C proteins species (Figure S1) (Ma *et al*, 2004; Dokland *et al*, 2004; Shang *et al*, 2018; Pulkkinen *et al*, 2018; Poonsiri *et al*, 2019; Barrass *et al*, 2021). The TBEV C protein is a dimer both in solution, and within the virion, although higher-order oligomerisation has been suggested to occur in other flaviviruses such as Zika virus (Kiermayr *et al*, 2004; Kaufman *et al*, 2020; Tan *et al*, 2020; Barrass *et al*, 2021). The C protein contains a positively-charged, flexible N-terminus (17 aa) and four α-helices (α_1_–α_4_), with dimerization occurring via antiparallel interactions of the α_2_ and α_4_ helices (Ma *et al*, 2004; Dokland *et al*, 2004; Shang *et al*, 2018; Boon *et al*, 2018; Pulkkinen *et al*, 2018; Poonsiri *et al*, 2019; Barrass *et al*, 2021). The α_4_ helix contains multiple positively-charged residues, thought to form a charged surface responsible for nucleic acid binding (Samuel *et al*, 2016; Shang *et al*, 2018; Kaufman *et al*, 2020). The other side of the dimer (the α_2_ helix or the N-terminus) may be important for nucleic acid or membrane binding, although this facet of C protein function remains under-studied (Markoff *et al*, 1997; Kofler *et al*, 2002; Nemésio *et al*, 2013; Shang *et al*, 2018).

As the C protein has membrane affinity, and the assembly process occurs at the ER, it is likely that specific ER lipids are involved in the process. Budding involves generating extreme membrane curvature which is only possible in the presence of suitable lipids. Flavivirus infection leads to a significant increase in curvature-promoting lipids in the host, and increases the curvature of the ER membrane (Miller *et al*, 2007; Gillespie *et al*, 2010; Perera *et al*, 2012; Peña & Harris, 2012; Martín-Acebes *et al*, 2014; McMahon & Boucrot, 2015; Cortese *et al*, 2017; Yau *et al*, 2019). Previous work has mainly concentrated on fluorescence microscopy-based characterization of C protein localization in infected cells, experiments conducted with C-derived peptides, and indirect characterization of membrane insertion (Markoff *et al*, 1997; Kofler *et al*, 2002; Samsa *et al*, 2009; Nemésio *et al*, 2011, 2013; Fajardo-Sánchez *et al*, 2017; Vonderstein *et al*, 2018). Here, we have probed the role of lipids in the NC assembly by using recombinant TBEV C and various model lipid systems to biophysically characterize the interaction of the C protein with membranes. We show that the C protein initially binds to lipids with negatively-charged head groups via electrostatic interactions which is then followed by membrane insertion. We confirm the physiological relevance of the experimental system by showing that the C protein can recruit genomic RNA whilst membrane-bound in the presence or absence of the 17 N-terminal residues. Furthermore, we characterise the TBEV lipidome with ultra-high performance liquid chromatography-quadrupole time-of-flight mass spectrometry (UHPLC-QTOF-MS) and show that the TBEV virion contain lipids with negatively-charged head groups. Our results indicate that the C protein may be recruited to the ER via positively charged lipid head groups, helping to nucleate the assembly of the NC and acquisition of the membrane through budding into the ER.

## Results

### Purified C protein binds to lipids with negatively-charged head groups

Both the full-length C and the truncated C (C_18–93_, containing residues 18–93) were expressed in *E.coli*, and after immobilized metal affinity chromatography and ion-exchange chromatography, eluted as pure dimers in size-exclusion chromatography confirmed by SDS-PAGE and immunoblotting (Figure S2). The A_260_/A_230_ ratios of the purified proteins were 0.61±0.02 and 0.55±0.25 for C and C_18–93_ respectively indicating no significant nucleic acid contamination (average of three technical replicates) (Warburg & Christian, 1942).

To determine if C protein bound to membranes and if lipid binding was dependent on the type of lipid head groups, we used liposome co-sedimentation. First, we used liposomes containing either brain total lipid extract (BTLE), or a mixture of the uncharged lipids 1,2-dioleoyl-sn-glycero-3-phosphocholine (DOPC) and 1,2-dioleoyl-sn-glycero-3-phosphoethanolamine (DOPE). C protein was found to bind and co-sediment with BTLE liposomes, but not with DOPC-DOPE (Figure 1A, B). Next, we supplemented the DOPC-DOPE liposomes with individual glycerophospholipids, sphingolipids or cholesterol (Chol) representing different specific lipid properties. Incorporation of sphingomyelin (SM), phosphatidylinositol (PI), or galactocerebrosides (GalCer) showed that lipids with uncharged or glycosylated head groups did not markedly increase C protein binding, neither did Chol. However, clear binding was detected after incorporation of the negatively-charged 1-palmitoyl-2-oleoyl-sn-glycero-3-phospho-L-serine (POPS) or the highly negatively-charged L-α-phosphatidylinositol-4,5-bisphosphate (PI(4,5)P_2_) (Figure 1A, B). This indicated that C bound to negatively-charged membranes without a preference for a specific head group. Furthermore, PI(4,5)P_2_ that is 3 times more charged than POPS resulted in increased binding (Figure 1A, B). To further confirm the role of charge, DOPC-DOPE liposomes were supplemented with increasing concentrations of POPS, increasing the negative charge of the liposomes proportionally, and co-sedimentation was then assessed. The fraction of co-sedimenting C protein increased with increasing POPS concentrations, with all C protein being bound at 40 % POPS (Figure 1C, D). Thus, 40 % POPS was used for further experiments.

**Figure 1.**
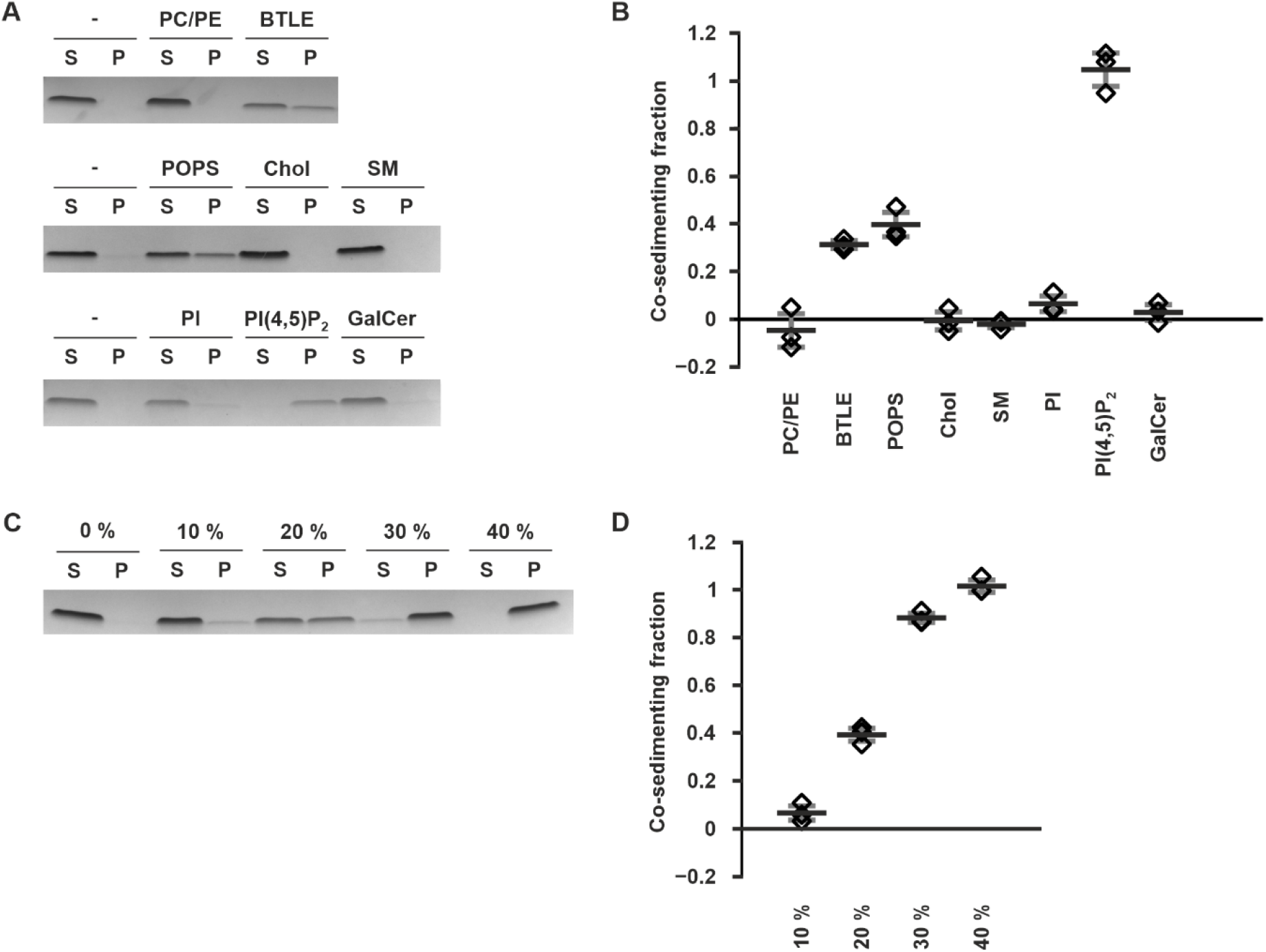
C protein binds preferentially to negatively-charged lipids **A, B** C protein co-sedimentation with liposomes of different compositions. (**A**) shows representative SDS-PAGE bands with S indicating the supernatant and P the pellet. (**B**) shows densitometric quantitation of the co-sedimenting fraction from the pellet. PC/PE refers to pure DOPC/DOPE liposomes, BTLE to pure BTLE liposomes, a dash to the no liposome control and the other labels to DOPC/DOPE liposomes supplemented POPS, Chol, SM, PI, PI(4,5)P_2_ or GalCer. **C, D** C protein co-sedimentation with liposomes containing varying amounts of POPS. (**C**) shows representative SDS-PAGE bands where S indicates the supernatant and P the pellet with the POPS percentage indicated above the lanes. (**D**) shows densitometric quantitation of the co-sedimenting fraction from the pellet with the w/w percentage of POPS in the liposomes is indicated on the Y axis. Data information: In (**B**) and (**D**), data are shown as the averages of three replicates with the error bars representing the standard deviation (s.d.). Individual measurements are shown as diamonds. Data are normalized against no liposome (**B**) or 0 % POPS control (**D**).

### Initial C protein recruitment to the membrane is of an electrostatic nature

To further characterise the binding of C protein to membranes, supported lipid bilayers (SLBs) on a quartz crystal microbalance with dissipation monitoring (QCM-D) measurements were used. SLBs were generated either from pure 1-palmitoyl-2-oleoyl-sn-glycero-3-phosphocholine (POPC) or a mixture of POPC and POPS (60:40 % w/w) (Figure 2 A). QCM-D measures the change in frequency (ΔF) of an oscillating quartz crystal which is related to the mass adsorbed to its surface. It also monitors the energy dissipation (ΔD) representing how rigid the adsorbed material is, a high dissipation corresponding to a more viscoelastic (soft) film. During formation of SLBs the adsorption of liposomes and subsequent fusion into a two-dimensional rigid SLB can be scored using these parameters (Figure 2A) (Cho *et al*, 2010). After the formation of the SLB and the equilibration of the system with Tris-buffered saline (TBS), the ΔF and ΔD values were zeroed, and the C protein was injected. The C protein injection caused the ΔF to decrease to −23.9 Hz on average (s.d. 2.2 Hz) in POPC/POPS experiments indicating a clear binding of additional mass on to the SLB (Figure 2B, C). However, repeating the experiment in the presence of pure POPC SLBs, resulted in a significantly smaller ΔF after C protein injection (on average 0.9 Hz, s.d. 0.88 Hz) indicating no mass binding (Figures 2 C, S3). The ΔF difference between protein binding to the POPC/POPS and POPC SLBs was statistically significant (p-value 6.94×10^−7^). Together, this comparison showed that the binding of C protein to SLBs was dependent on the presence of charged POPS (Figures 2 C, S3). Interestingly, the ΔD upon C protein adsorption to the POPC/POPS SLBs was very low (3.8×10^−7^, s.d 1.4×10^−6^) indicating that the C protein forms a very rigid layer when adsorbed to the membrane (Cho *et al*, 2010).

**Figure 2.**
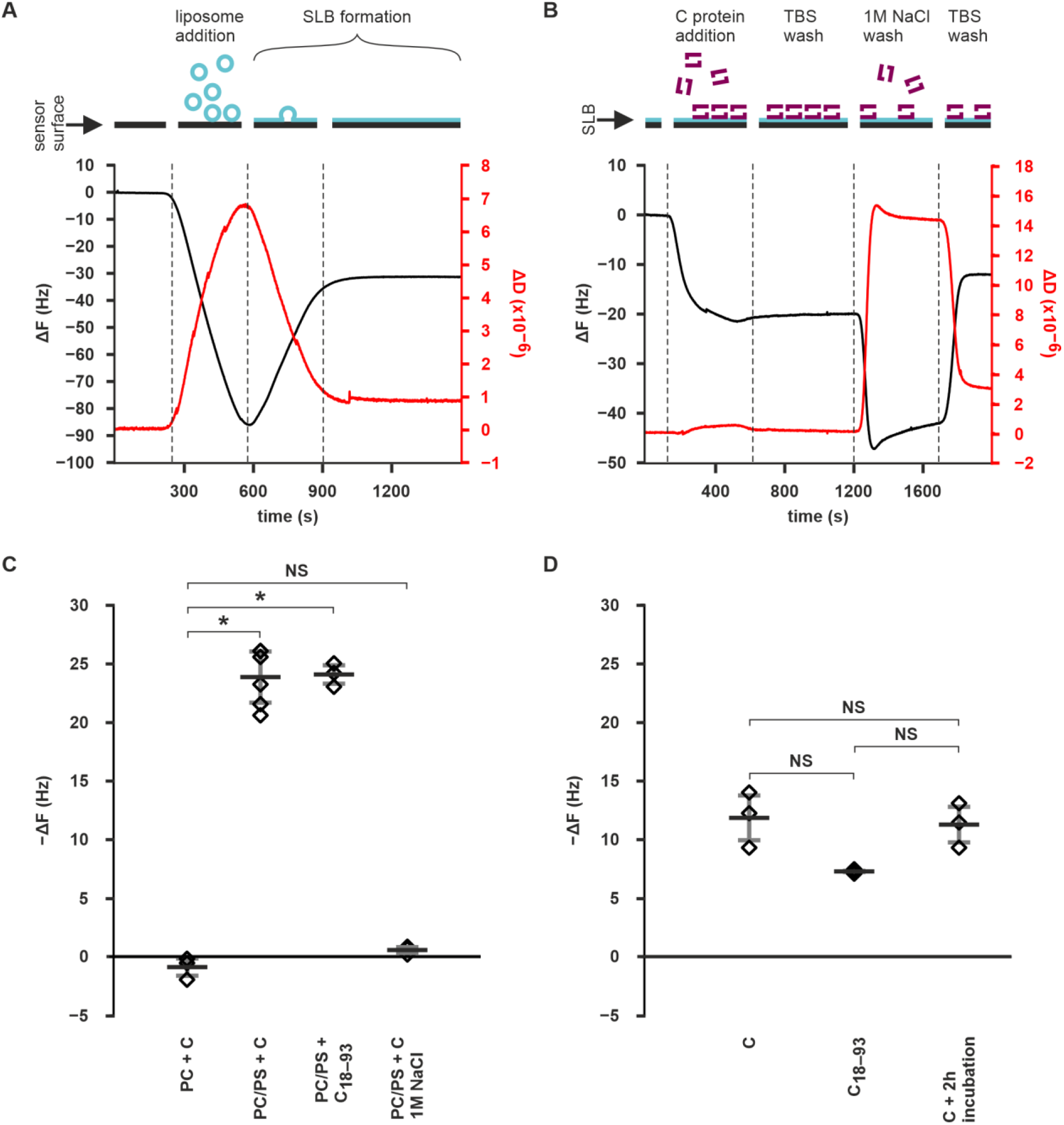
C protein binding to SLBs in QCM-D measurements **A** Representative QCM-D curves from POPC/POPS SLB generation. The ΔF and ΔD have been zeroed to prelipid injection equilibrium values. **B** Representative QCM-D curves from a C protein binding experiment on POPC/POPS SLB. The ΔF and ΔD have been zeroed to equilibrium values after SLB formation. The large reversible changes in ΔF and ΔD after 1 M NaCl injection indicate a typical “buffer shift”, as the ΔF and ΔD values are sensitive to buffer viscosity (Liu *et al*, 2022). **C** Quantitation of the final −ΔF values upon C protein and C_18–93_ injection on SLBs composed of pure POPC (PC), or POPC/POPS mixture (PC/PS) either in TBS or in 1 M NaCl-containing buffer. **D** Quantitation of the final −ΔF values on SLBs with prebound C or C_18–93_ after 1 M NaCl wash either right after equilibration, or after 2h incubation. Data information. Data in **C** and **D** are shown as the mean of 6 (C binding in TBS on POPC/POPS) or 3 (all others) replicates with the s.d. represented by the error bars. Individual measurements are shown as diamonds. Statistical significance (p < 0.05) is indicated by an asterisk and non-significant differences with NS. See main text for the p-values.

To probe the electrostatic nature of the initial C protein recruitment to the membrane, the C protein was injected in a buffer containing 1 M NaCl on SLBs preequilibrated with the same buffer. After reequilibrating the SLBs with TBS, the ΔF value was on average −0.6 Hz (s.d 0.3 Hz) compared to the pre 1 M NaCl equilibration showing that no C protein bound to the SLBs (Figures 2 C, S3). The difference between injection on pure POPC SLBs and on POPC/POPS SLBs in 1M NaCl was not statistically significant (p-value 0.13). To test if the C protein membrane contacts are purely electrostatic, C protein was bound to SLBs in TBS, and then washed with 1 M NaCl buffer either right after equilibration or after a 2 h post-equilibration incubation. A wash with 1M NaCl resulted in what is referred to as “buffer shift” in the F and D values, which was reversed when moving back to the original buffer (Liu *et al*, 2022). In both cases, only partial detachment of the C protein from the SLBs was detected with the final ΔF value being −11.9 Hz in the immediate wash experiment, and −11.3 in the 2h-incubation experiment (s.d. 1.9 and 1.5 Hz, respectively) (Figure 2 D). The ΔF values were significantly different from zero with p-values of 0.013 (immediate wash experiment) and 0.009 (2h-incubation experiment). However, they were not significantly different between the experiments with a p-value of 0.10. This shows that initial membrane recruitment of C protein is strongly dependent on its interactions with the negatively-charged lipid headgroups. However, once bound, the C protein-membrane interaction is complemented with non-electrostatic interactions such as membrane insertion or protein oligomerization. Since the amount of detached protein is not dependent on incubation time, it suggests that the protein’s non-electrostatic organization on the membrane occurs concurrently with adsorption.

When the same measurements were performed with the C_18–93_ construct, similar results to the wild-type C were obtained. The C_18–93_ protein injection caused a 24.1 Hz drop in ΔF (s.d 0.8 Hz) with the final ΔF value being significantly different from the C protein binding to pure POPC experiment (p-value 6.04×10^−7^) (Figure 2 C, S3). When the bound C_18–93_ was washed with high-salt buffer, mass was retained on the SLB with a final ΔF value of −7.3 Hz (s.d. 0.2 Hz) (Figure 2 D, S3). The final ΔF value was significantly different from zero (p-value 0.0002). Additionally, the final ΔF value was not significantly different from the full-length C protein experiments (p-values 0.079 and 0.066 for the comparison to the immediate wash and 2h incubation experiments respectively). This showed that the deletion of the N-terminus did not significantly alter membrane binding of the C-protein.

### C protein inserts into membranes

To investigate if the C protein membrane binding includes insertion into the membrane after the initial electrostatic binding, we used Langmuir-Blodgett trough monolayer experiments. We formed monolayers of pure POPC or POPC/POPS mixture (60:40 %, w/w) and monitored the pressure (π) of the monolayers following a C or C_18–93_ injection into the subphase. The π increase (Δπ) between the starting (π_0_) and ending pressures were measured at different π_0_ values. The regression between Δπ and π_0_ was used to determine the maximum injection pressure (MIP) of the proteins.

Injection of full-length C into the subphase led to sharp π increases at all tested π_0_ values in monolayers containing 40 % POPS (Figures 3A, S4). The MIP of full-length C was 39.1 mN/m, which is remarkably higher than the pressure of a lipid bilayer (~30 mN/m), suggesting that C is capable of inserting into bilayers such as the ER membrane (Figure 3B) (Blume, 1979; Marsh, 1996; Calvez *et al*, 2009). Contrastingly, in the absence of POPS, smaller π increases were detected, and the Δπ value was zero at π_0_ of 20.20 mN/m (Figures 3A, S4). Furthermore, the MIP value of full-length C in pure POPC monolayers was 19.7 mN/m, which is notably smaller than the pressure of a lipid bilayer showing that negatively-charged lipids such as POPS are needed for C protein membrane insertion at physiological membrane pressures (Figure 3C) (Blume, 1979; Marsh, 1996; Calvez *et al*, 2009). The C_18–93_ protein behaved similarly to the wild-type C in POPS-containing monolayers with a MIP of 37.7 mN/m (Figures 3A, D, S4). This corroborates the similar QCM-D results between C and C_18–93_, and shows that the N-terminus is not needed for C protein membrane insertion.

**Figure 3.**
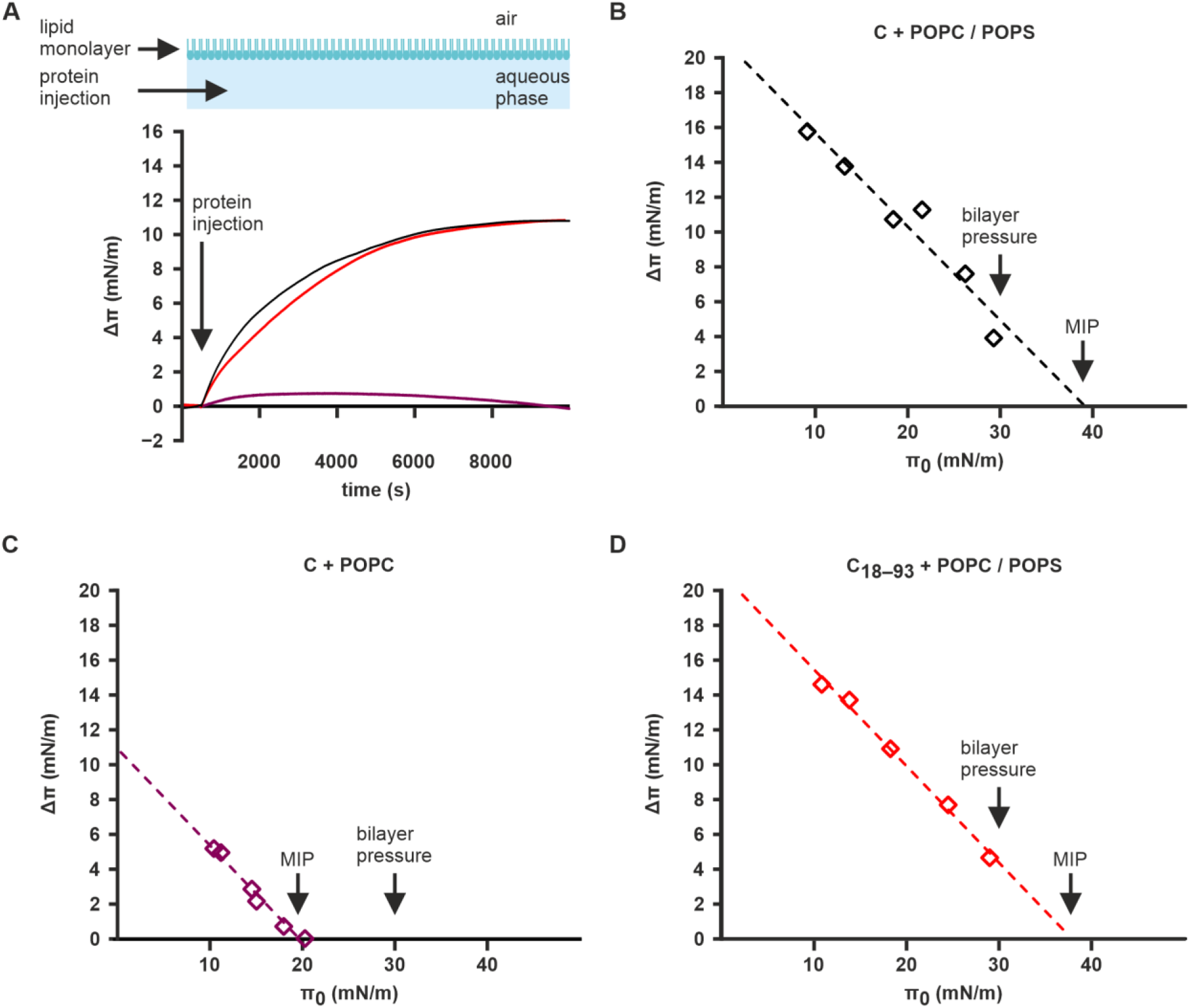
C protein membrane insertion **A** Representative Δπ curves for C protein injected with a POPC/POPS (black) or POPC (purple) monolayer, and C_18– 93_ protein injected in with a POPC/POPS monolayer (red) at π_0_ values of 18.1±0.2 mN/m. **B, C, D** MIP curves of C protein with POPC/POPS (**B**) and pure POPC (**C**) monolayers, and C_18–93_ with POPC/POPS monolayers (**D**). The pressure of a bilayer (~30 mN / m) is indicated along with the MIP (Blume, 1979; Marsh, 1996; Calvez *et al*, 2009). Data information. In (**A**), π is zeroed to π_0_. The Individual measurements in (**B**), (**C**), and (**D**) are shown as diamonds. The R^2^ values in (**B**), (**C**), and (**D**) are 0.9344, 0.9818, and 0.9935, respectively.

### Membrane-bound C protein can recruit TBEV genomic RNA

To confirm that the C protein is biologically active, we investigated its ability to bind RNA. First, we confirmed the binding in solution using *in vitro* transcribed TBEV genomic RNA with a gel electrophoretic mobility shift assay (GEMSA). Using a C to RNA molar ratio of 624:1 fully prevented the RNA from migrating normally indicating binding (Figure 4A).

**Figure 4.**
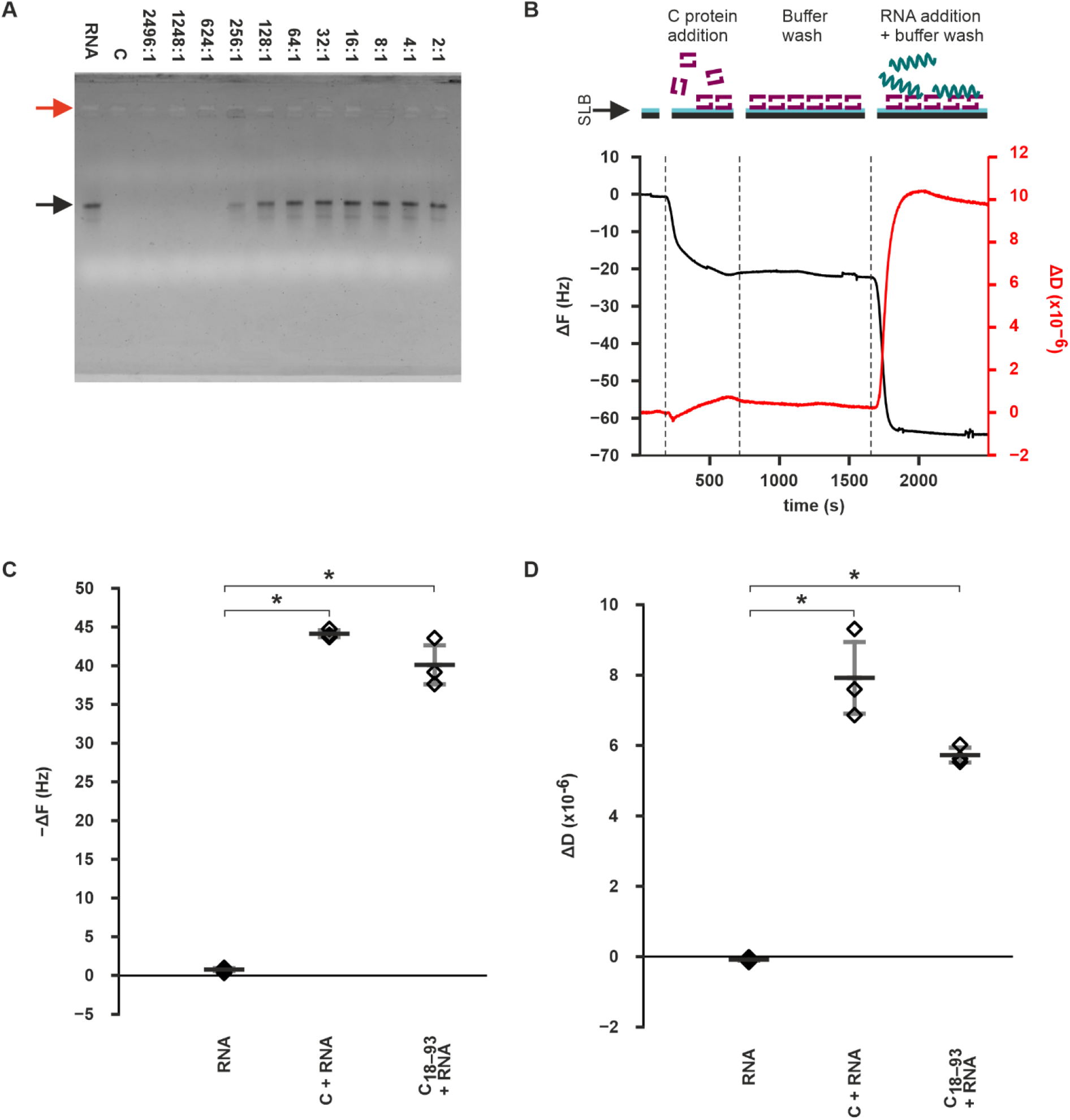
C protein - RNA interactions **A** GEMSA gel showing C protein binding to RNA in solution. RNA refers to only RNA, and C to only C protein corresponding to the highest C protein concentration used. The C to RNA molar ratio is indicated. The red arrow indicates the wells where the C-RNA complex has been immobilized, and the black arrow indicates the bands of the freely migrating RNA. **B** Representative QCM-D curves from an RNA-binding experiment on POPC/POPS SLBs with bound C protein. The ΔF and ΔD have been zeroed to equilibrium values after SLB formation. **C, D** Quantitation of ΔF (**C**) and ΔD (**D**) changes upon RNA insertion on SLBs pretreated with C, C_18–93_, or no protein. Data information. Data in C and D are shown as the mean of 3 repeats with s.d. represented by the error bars. Individual measurement are represented by the diamonds. Statistical significance (p < 0.05) is indicated with an asterisk. See the main text for the p-values.

As the simultaneous binding of membranes and RNA is a key function of the C protein, we investigated its RNA recruitment capability whilst membrane-bound using QCM-D. In these experiments, C protein was essential for the recruitment of TBEV genomic RNA on SLBs (Figure 4B, C, D). Injecting 5 μg of *in vitro* transcribed TBEV genomic RNA on SLBs which had bound C protein resulted in an average ΔF of −44.2 Hz at equilibrium (s.d. 0.5 Hz) showing RNA binding (Figure 4B, C). In contrast, the ΔF value with SLBs with no prebound C remained close to zero after RNA injection, with an average equilibrium ΔF of −0.6 Hz (s.d. 0.3 Hz) indicating no binding of mass on the SLBs (Figures 4C, S5). The equilibrium ΔF value difference between C protein-containing and protein-free SLBs was statistically significant (p-value 1.34×10^−6^) (Figure 4C). Interestingly, the ΔD value also increased to 7.92×10^−6^ on average after RNA binding (s.d. 1.03×10^−6^) in the protein-treated SLBs, indicating that the RNA did not bind as a rigid uniform layer but rather as a flexible assembly (Figure 4B, C). Without prebound C protein, the ΔD did not increase and remained at an average −8.8×10^−8^ at equilibrium (s.d. 5.4×10^−8^) (Figures 4C, 5). The equilibrium ΔD value difference between the experiments with and without prebound C protein was statistically significant (p-value 0.008) (Figure 4D). These data show that membrane-bound C protein is capable of recruiting TBEV genomic RNA at the membrane, suggesting that this also happens in the context of NC assembly.

When the SLB RNA-binding experiments were repeated with C_18–93_, the results were similar as with full-length C. After RNA injection on C_18–93_-treated SLBs, the ΔF value dropped to −40.1 Hz on average and the ΔD value increased to an average of 5.7×10^−6^ (s.d. 2.5 Hz and 2.1×10^−7^, respectively). The difference to the no protein experiment for both ΔF and ΔD was significant (p-values 0.002 and 0.0007, respectively) (Figure 4C, D). While the ΔF and ΔD values in the C_18–93_ experiments were closer to zero than in the wild-type C experiments, the differences were not statistically significant (p-values 0.156 and 0.097 respectively).

### TBEV virions contain negatively-charged phosphatidylserine

If the binding of C protein to negatively-charged lipid head groups is important for particle assembly, it is likely that these lipid species are incorporated into the virion during the budding into the ER. To investigate this possibility, we used UHPLC-QTOF-MS to characterize the TBEV lipid content of purified virions. The average integrated peak area of each investigated lipid species was compared between TBEV-containing samples, and buffer-only mock samples. Significantly higher integrated peak areas were interpreted as the presence of the species in the TBEV preparation, instead of noise caused e.g. by buffer or instrument contamination.

Multiple lipid species of different classes were detected in the TBEV samples (Figure 5 A, file S1). The virions contained lipids from two negatively-charged classes, phosphatidylserine (PS) and phosphatidylglycerol (PG) (Figure 5 A, File S1). In addition, neutral carnitine fatty acids (Car), phosphatidylethanolamines (PE), PIs, phosphatidylcholines (PC), ceramides (Cer), hexosylceramides (HexCer), and triglycerides (TG) were detected (Figure 5 A, File S1). The analysed lipid classes not found in virions were lyso-PE (LPE), lyso-PC (LPC), diglyceride (DG), and cardiolipin (CL) (Figure 5B, File S1). The virions contained multiple PS species: 32:1, 34:1, 36:1, 36:2, and 38:5 (number of carbons:number of double bonds), which confirms the biological relevance of using this lipid class in the model membrane experiments (Figure 5B, File S1). These data show that negatively-charged lipids are incorporated into the virion during the budding process, supporting the hypothesis that C protein interactions with them have a role in NC assembly and budding.

**Figure 5.**
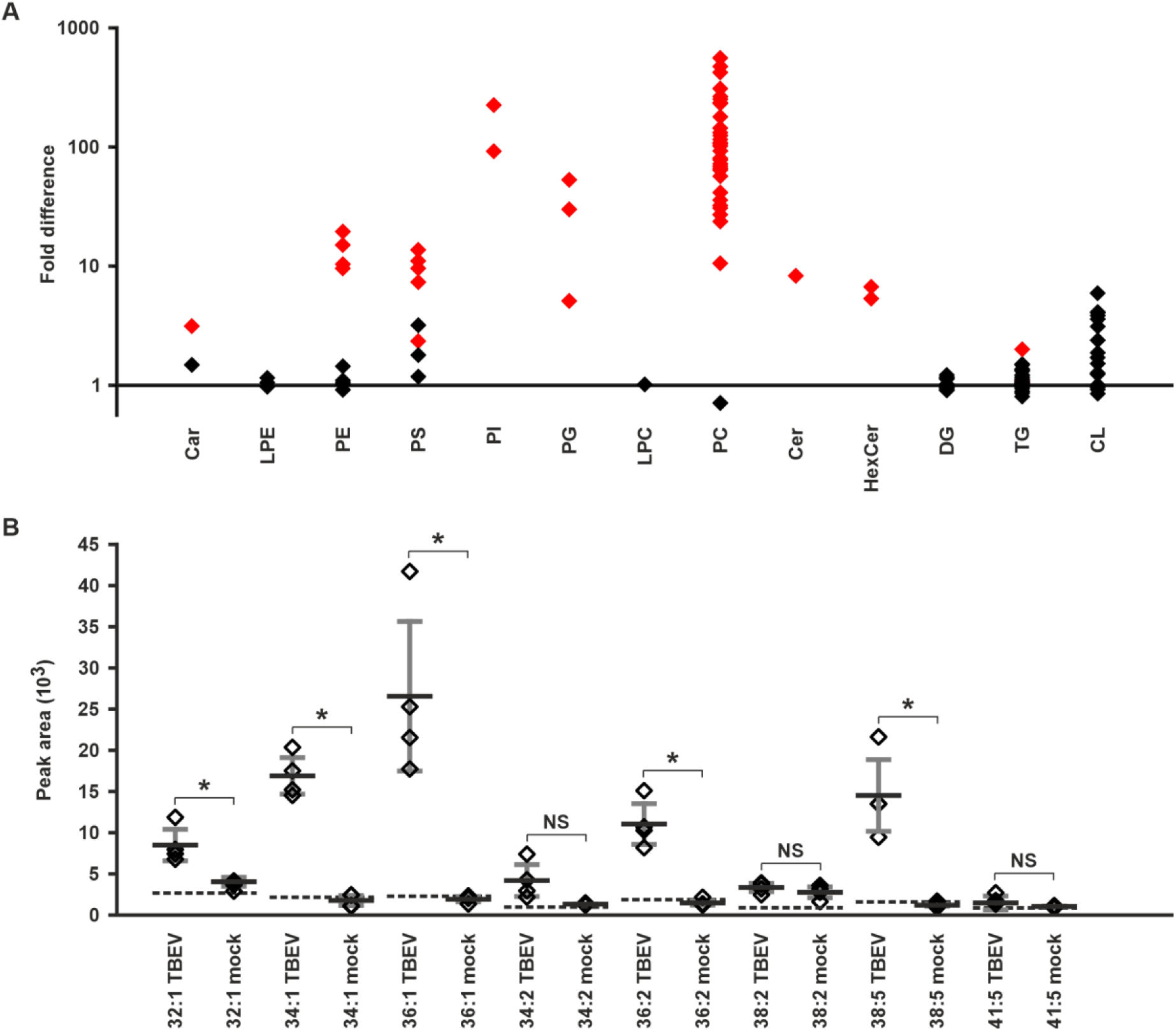
Mass spectrometry characterization of the lipids in TBEV virions **A** Fold-increases of the integrated peak areas of the analysed lipid species in TBEV versus buffer samples. **B** Integrated peak areas of the detected PS species in TBEV virions and mock samples. The background peak level of each lipid species from a blank run is indicated with the dotted line. Data information: Data in **A** are shown as the ratio of the average integrated peak areas of TBEV virions divided by the buffer controls from four technical repeats. Data in **B** are shown as the averages of four technical replicates with the horizontal bar representing the mean and the error bars representing the s.d.. Individual species (**A**) or measurements (**B**) are represented by the diamonds. Significance in **A**, and **B** is indicated when p < 0.05 either in red (A) or with an asterisk (B). For the p-values, see file S1.

## Discussion

Our unique combination of biophysical methods and model membrane systems allowed us to show that negatively-charged lipids are required for TBEV C protein membrane interactions, and that the protein inserts into membranes. The methods allowed us to demonstrate RNA recruitment onto membranes by the C protein, supporting that the NC assembly occurs on the ER membrane (Pulkkinen *et al*, 2018). We show that aa residues 1–17 are not required for these interactions, and that membrane binding and RNA recruitment can occur independently of any other host or viral proteins. Additionally, we confirm the biological relevance of the interactions with negatively-charged lipids observed in the biophysical experiments by showing that negatively-charged PS and PG lipids are incorporated into the virion. PS, not PG, has been found previously in the lipid composition of West Nile virus (Martín-Acebes *et al*, 2014). Our findings on C protein membrane binding properties agree with previous observations from flaviviruses (Markoff *et al*, 1997; Kofler *et al*, 2002; Nemésio *et al*, 2011; Freire *et al*, 2013; Nemésio *et al*, 2013; Ambroggio *et al*, 2021).

While it is clear that the C protein can perform both membrane binding and insertion as well as RNA binding simultaneously, the mechanism for this remains unclear. The first 17 residues of the C have been reported to be dispensable for TBEV particle assembly, which is consistent with our *in vitro* results, and supports that the N-terminus primarily has a role in modulating the host response (Kofler *et al*, 2002; Yang *et al*, 2002; Limjindaporn *et al*, 2007; Colpitts *et al*, 2011; Bhuvanakantham & Ng, 2013; Katoh *et al*, 2013; Urbanowski & Hobman, 2013; Samuel *et al*, 2016; Slomnicki *et al*, 2017; Fontaine *et al*, 2018). As the initial membrane binding and RNA recruitment are charge-mediated and the charged N-terminus is not needed for either of these functions, it seems like the positively-charged α_4_–α_4_ interface would bind both, unless conformational changes occur during membrane insertion expose additional binding sites (Figure S1) (Pulkkinen *et al*, 2018; Barrass *et al*, 2021). Based on the QCM-D results, the binding followed two different kinetics, and after binding and insertion, only half of the C protein could be removed by a high NaCl wash. This indicates that most probably, higher-order interactions occur, with rapid nucleation on the membrane, for instance by the α_4_–α_4_ interface, followed by slower recruitment of additional molecules via the opposite face of the dimer, namely the α_2_ helix (Figures 2B and 4B). The newly added molecules would have the α_4_–α_4_ interface exposed for RNA recruitment. These additional molecules would more likely be removed by the high NaCl wash, as they would not be inserted into the membrane due to steric hindrance from the first layer of protein. The only evidence for regular interactions in the virion is that of a dimer from cross-linking mass spectrometry, but in order to form the NC, it is highly likely that there are additional, higher-order C protein interactions (Barrass *et al*, 2021). The results support that C binding to negatively-charged surfaces is an important step in RNA recruitment, just as in dengue virus (DENV) (Mebus-Antunes *et al*, 2022). We cannot rule out that the C protein preparation is heterogeneous, despite its appearance, and this heterogeneity could cause the partial insertion observed. Further functional studies are needed to determine the orientation, conformation and multimerisation status of C on the membrane.

Another factor that could influence the C protein interaction with negatively-charged lipids, would be its phosphorylation state. In West Nile virus, C protein, phosphorylated by protein kinase C, is imported into the nucleus through binding to importin α. The nuclear import leads to localization in the nucleolus where C binds to HDM2 starting apoptosis (Bhuvanakantham *et al*, 2010). Phosphorylation decreases the cytoplasmic C protein amount, and reduces binding to RNA (Cheong and Ng, 2011), thus decreasing the pool of protein available for NC assembly. TBEV C protein is also transported to the nucleolus (Vonderstein *et al*, 2018), so it is likely that this is also through phosphorylation and importin α. Likely residues for phosphorylation are S83, S19 and S65. One of the predicted phosphorylation sites (S83) is present on the α_4_ helix, so the charge of the phosphate group could repel the negatively-charged membrane and prevent binding, whereas dephosphorylation would promote assembly, increasing the propensity of C to bind to both membranes and RNA in TBEV. Such a mechanism has recently been reported for herpesvirus nuclear egress complexes that bind to negatively-charged lipids (Thorsen *et al*, 2021).

Based on our results, negatively-charged ER lipid patches may act as the nucleation sites for NC assembly by binding the C protein that subsequently recruits the genomic RNA. PS, which we detected in the virion envelope, is also a major component of the ER (Van Meer *et al*, 2008). ER lipid microdomains are most prevalent at the contact sites between the ER, mitochondria (rich in PG), and lipid droplets where there is extensive exchange of lipids between organelles (Van Meer *et al*, 2008; Wang *et al*, 2020). Mitochondria and lipid droplets are often observed in close proximity to the ER replication sites of TBEV, DENV and Zika virus (Chatel-Chaix *et al*, 2016; Cortese *et al*, 2017; Yau *et al*, 2019). Additionally, the recruitment of the C protein to lipid droplets has been suggested to be important for DENV assembly (Samsa *et al*, 2009; Carvalho *et al*, 2012). As flavivirus infections cause changes in the host cell lipid metabolism, it is possible that this leads to an increase in PS and PG content in the ER. The observed frequent association of mitochondria and lipid droplets with flavivirus replication sites as well as the presence of PS and PG in the TBEV envelope maybe the results of this (Perera *et al*, 2012; Peña & Harris, 2012; Martín-Acebes *et al*, 2014).

In conclusion, we have determined that NC assembly is most likely membrane-associated, involving specific lipids. Although often ignored in virus structure and assembly, the importance of specific lipid species is gradually being recognised in flavivirus assembly, including the recent discoveries of ordered lipids in flavivirus membranes coordinated by the E and M membrane helices (DiNunno *et al*, 2020; Khare *et al*, 2021; Hardy *et al*, 2021; Renner *et al*, 2021; Pulkkinen *et al*, 2022). Our results provide a starting point for further structural and mechanistic studies on the role of lipid interactions in TBEV NC assembly.

## Materials and methods

### Lipids

The following lipids were purchased from Avanti Polar Lipids (Alabama, USA) as lyophilized powder: DOPC (850375), DOPE (850725), porcine BTLE (131101), POPS (sodium salt, 840034), SM (N-[dodecanoyl]-sphing-4-enine-1-phosphocholine, LM2312), ovine Chol (700000), bovine liver L-α-PI (840042), porcine brain PI(4,5)P_2_ (ammonium salt, 840046), and POPC (850457). Bovine brain GalCer (C4905) were purchased from Sigma-Aldrich (Missouri, USA) as lyophilized powder.

### Cloning and protein purification

Full-length TBEV C protein (residues 1–96) and a truncated C protein (C_18–93_, containing residues 18–93) from the Kuutsalo-14 strain (GenBank: MG589938.1) with an N-terminal 6x His + Small Ubiquitin-like Modifier (SUMO) tag were cloned into pET28 plasmid as a commercial service by DNA Dream Lab (Helsinki, Finland) (Kuivanen *et al*, 2018). The proteins were expressed in *E. coli* ArcticExpress cells. The cells were grown o/n at 37 °C with 225 RPM shaking in lysogeny broth (LB, 1% w/v tryptone, 0.5% w/v yeast extract, 85 mM NaCl) containing 50 μg/ml kanamycin and 10 μg/ml gentamycin. Cells were inoculated into fresh LB without antibiotics, and grown at 30 °C with 225 RPM shaking until OD600 reached 0.5. The cultures were moved to +12 °C with 225 RPM shaking for an hour and induced with isopropyl β-d-1-thiogalactopyranoside (IPTG, final concentration 1 mM). The induced cultures were grown for 24 h at 12 °C with 225 RPM shaking and harvested by centrifuging at 3000 × *g* for 20 min at +4 °C. The supernatant was discarded, and the pellets stored at −80 °C until needed.

The frozen pellets were resuspended on ice in 300 mM NaCl, 50 mM Tris, 375 mM L-arginine, 20 mM imidazole, pH 7.5 containing Pierce protease inhibitors (Thermo Fisher, A32965). The cells were lysed with an Emulsiflex apparatus (Avestin Inc., Ontario, Canada) operated at 15 000 psi for 10 min. The cell lysates were precleared for 30 min at 20 200 × *g* at + 4 °C. The resulting supernatant was further cleared by centrifuging at 96 600 ×*g* for 1.5 h at +4 °C. The proteins were captured with nickel immobilized metal affinity chromatography using a 5 ml HisTrap FF column (Cytiva, Massachusets, USA) and a linear imidazole gradient from 0 mM to 1000 mM over 25 column volumes. The purified proteins were buffer-exchanged into 300 mM NaCl, 50 mM Tris, pH 7.5 with PD 10 columns (Merck, New Jersey, USA) using the manufacturer’s protocol. The SUMO tags were cleaved using in-house-produced SUMO protease o/n (full-length C) or for 48 h (C_18–93_) at RT with gentle agitation. The cleaved tags were removed by cation exchange chromatography using a 1 ml HiTrap SP HP column (Cytiva) with a linear NaCl gradient from 300 mM to 1000 mM over 25 column volumes. SEC was used to buffer exchange the samples into 300 mM NaCl, 50 mM Tris, pH 7.5 using a Superdex 200 10/300 GL column (Cytiva). The peak fractions were collected, aliquoted, and stored in −80 °C. The aliquots were characterized with SDS-PAGE and immunoblotting (see below), the protein concentrations were measured with a QFX fluorometer (DeNovix, Delaware, USA) using a Qubit Flex kit (Thermo Fisher, Massachusetts, USA), and the A_260_/A_280_ ratios of the samples were measured using a Nanodrop instrument (Thermo Fisher).

### SDS-PAGE and immunoblotting

Samples were mixed with Laemmli sample buffer, incubated for 5 min at 95 °C, and proteins were resolved using electrophoresis in 4–20 % polyacrylamide gels (Bio-Rad, California, USA). Proteins were visualized using Coomassie blue staining or by immunoblotting. For immunoblotting, the proteins were transferred onto nitrocellulose membranes (GE Healthcare, Illinois, USA) using a Trans Blot turbo apparatus (BioRad) using the manufacturer’s protocol. The membranes were blocked using 5 % (w/v) milk in 150 mM NaCl, 20 mM Tris, 0.1 % TWEEN 20, pH 7.6 (TBST) for 30 min at RT with gentle rocking and washed with TBST. Membranes were incubated with the primary anti-C antibody (1:1000 dilution in TBST containing 5% milk) (Panayiotou *et al*, 2018). After incubation, the membranes were washed 3 times with TBST for five minutes with gentle rocking. The secondary antibody was diluted 1:10 000 in TBST, added on the membranes, and the membranes were incubated for 30 minutes at RT with gentle rocking. The secondary antibody used was IRDye 680 RD goat anti-rabbit IgG (Li-COR). After incubation, the membranes were washed with TBST and visualised using an Odyssey imager (Li-COR).

### Liposome sedimentation assay

The protocol was adapted from Larsson *et al*, 2020. Liposomes of various lipid species (see table 1) were prepared by diluting total of 0.5 mg of lipids in a 10:3 mixture of chloroform and methanol in glass tubes. Lipid films were prepared by drying the lipids under a gentle nitrogen stream while rotating the glass tubes. The lipids were further dried under a gentle nitrogen stream for 30 min. The lipids were rehydrated to 1 mg/ml by adding 500 μl 150 mM NaCl, 50 mM Tris, pH 7.6 (TBS) and by incubating at RT for 30 min. The liposomes were sonicated with a bath sonicator for 20 s (Transsonic T310, Elma Schmidbauer, Singen, Germany) and either used immediately, or stored at +4 °C for up to two days.

**Table 1.** Liposome compositions used for sedimentation assays

| Lipid species | composition (% w/w) |
| --- | --- |
| DOPC/DOPE | 60:40 |
| DOPC/DOPE | 50:50 |
| BTLE | 100 |
| DOPC/DOPE/POPS | 45:45:10 |
| DOPC/DOPE/POPS | 40:40:20 |
| DOPC/DOPE/POPS | 35:35:30 |
| DOPC/DOPE/POPS | 30:30:40 |
| DOPC/DOPE/Chol | 40:40:20 |
| DOPC/DOPE/SM | 40:40:20 |
| DOPC/DOPE/PI | 40:40:20 |
| DOPC/DOPE/PI(4,5)P<sub>2</sub> | 55:40:5 |
| DOPC/DOPE/GalCer | 40:40:20 |

Fresh C protein aliquots were thawed on ice and centrifuged for 10 min at 20 000 × *g* at + 4 °C, and the supernatant moved to fresh tubes. The protein was incubated either with experimental liposomes or TBS for 5 min at RT with a final protein concentration of 7 μM and a final lipid concentration of 500 μg/ml. The liposomes were pelleted by centrifuging for 20 min at 100 000 × *g* at RT. The supernatants were collected, and the pellets resuspended in 50 μl TBS. Both the supernatants and the pellets were analysed with SDS-PAGE as above, and the fraction of C protein co-sedimenting with the liposomes was quantified using Fiji (Schindelin *et al*, 2012).

### RNA synthesis and electrophoretic mobility shift assay

Full-length TBEV genomic RNA was produced from an infectious clone based on the TBEV 93/783 sequence (GenBank: MT581212) using *in vitro* transcription. Linearized DNA containing the clone was transcribed using the mMESSAGE mMACHINE SP6 Transcription kit (Thermo Fisher) according to the manufacturer’s protocol. RNA was purified by LiCl precipitation according to the transcription kit protocol. RNA concentration was measured using a nanophotometer (DeNovix).

Transcribed RNA (300 ng) was mixed with C protein at molar ratios ranging from 1:2 to 1:2496 in 150 mM NaCl, 50 mM Tris, pH 7.5 prepared with RNAse-free reagents (RNA buffer) and incubated on ice for 20 min. The samples were mixed with loading dye (final composition: 60 % v/v formamide, 0.0133 % w/v bromophenol blue, 0.0053 % v/v xylene cyanol, 0.72 % w/v Tris, 0.37 % w/v boric acid, 6.6 mM EDTA) and analysed with electrophoresis on a 0.5 % agarose gel containing SYBR Green I (Thermo Fisher) for 50 min at 70 V.

### SLBs and QCM-D

The protocol was adapted from Liu *et al*, 2022. Liposomes containing either pure POPC, or a mixture of POPC and POPS (60:40 % w/w) were prepared as above in 20 mM citric acid, 50 mM KCl, 0.1 mM EDTA, pH 4.5 (CKB). The liposomes were homogenized by extruding 11 times through a 50 nm polycarbonate filter (Nuclepore Track-Etched Membranes, Whatman, Maidstone, UK) using a Mini Extruder instrument (Avanti Polar Lipids). QCM-D experiments were conducted with an X4 instrument equipped with a flow chamber (AWSensors, Valencia, Spain) using wrapped 14 mm 5 MHz, Cr/Au-SiO_2_ polished QCM-D sensors (AWSensors) at +23 °C. The sensors were incubated overnight in 2% SDS and cleaned with a UV-ozone cleaner (Bioforce Nanosciences, Iowa, USA) for 30 min before use. Frequency and dissipation were monitored for overtone 3, although they were measured for overtones 1, 5, 7, 9, and 11 as well. The chambers were filled with CKB, and the SLBs were formed by injecting 150 μl of the liposomes at a concentration of 0.1 mg/ml. After the formation of the SLBs, TBS was injected until the system reached equilibrium. For the 100% POPC SLBs, 7.7 μg of C protein in 100 μl TBS (corresponding to a concentration of 7 μM) was injected. For the 60 % POPC/40 % POPS SLBs, full-length C or C_18–93_ was injected either as above or the SLBs were washed with 1000 mM NaCl, 50 mM Tris, pH 7.6 (HSQB) and C protein injected as above but in HSQB. After C protein injection, the chambers were washed with TBS until equilibrium. In the experiments where C protein was injected in TBS, the protein-containing SLBs were washed with HSQB either immediately after reaching equilibrium, or after 2h incubation. Similarly, in C_18–93_ experiments, the protein-containing SLBs were washed with HSQB immediately after equilibrium was reached. After HSQ wash, the SLBs were washed with TBS until equilibrium. For the RNA experiments, the SLBs were washed with RNA buffer either without protein addition, or after an injection of C or C_18–93_ as above. 5 μg of *in vitro* transcribed TBEV genomic RNA in 100 μl RNA buffer was injected, followed with a wash with RNA buffer, and a wash with TBS until equilibrium. The equilibrium ΔF and ΔD values for each step were calculated as the average of a 30 s period starting 400 s after the start of the final TBS wash.

### Lipid monolayer experiments

The protocol was adapted from (Liu *et al*, 2022). Lipid formulations of either pure POPC or a 60:40 % (w/w) mixture of POPC and POPS were prepared in chloroform (final lipid concentration 0.5 mg / ml). A Microtrough G1 system (Ø 53 mm x 4 mm, Kibron, Helsinki, Finland) was used for the experiments. The trough was covered to prevent evaporation, and the experiments were conducted at +21 °C. The subphase was formed of TBS and constantly stirred with a magnetic stirrer during the measurement. π was measured with a Wilhelmy paper plate presoaked in TBS and calibrated for each measurement. The monolayers were formed by adding the lipid solutions on to the subphase in a dropwise manner and letting the chloroform evaporate. Once the initial π_0_ value of the experiment was reached, the monolayer was left to equilibrate for 30 min. Then, C protein or C_18–93_ in TBS was injected with a syringe under the monolayer via an injection port to reach the final protein concentration of 30 nM in the subphase. The π value was monitored until it plateaued. The π_0_ values were calculated as an average of the 30 s preceding the injection, and the final π values were calculated as the average of the 30 s surrounding the highest value reached. The MIP-values were determined by plotting Δπ against π_0_ and extrapolating the Δπ/π_0_ plot to the x axis.

### Cell Culture

Human neuroblastoma SK-N-SH cells (gift from Prof. Olli Vapalahti, University of Helsinki) were maintained in low-glucose Dulbecco′s Modified Eagle′s Medium (DMEM, Sigma) supplemented with 10 % foetal bovine serum (FBS, Gibco), 0.5 mg/ml penicillin, 500 U/ml streptomycin (Lonza Bioscience, Hayward, CA, USA), and 2 mM glutamine (Gibco, Waltham, MA, USA). The cells were maintained at +37 °C in a 5% CO_2_ atmosphere.

### Virus Propagation and Titration

TBEV-Eu strain Kuutsalo-14 passage 1 (gift from Prof. Vapalahti) was used to infect confluent SK-N-SH cells using a multiplicity of infection of 0.003 (Kuivanen *et al*, 2018). The cells were washed twice with phosphate-buffered saline (PBS), virus was added in infection medium (low-glucose DMEM, 0.5 mg/ml penicillin, 500 U/ml streptomycin, 2 mM glutamine, 2% FBS, 35 nM rapamycin [Bílý *et al*, 2015]), and incubated at +37 °C in 5% CO_2_. At 72 h post-infection, the supernatant was collected, and centrifuged for 10 min at +4 °C at 3800 × g. After centrifugation, the supernatant was collected and immediately purified (see below), or titered and stored at −80 °C.

For titration, virus samples were serially diluted 10 fold in infection medium. Confluent SK-N-SH cells in 6-well plates were washed twice with PBS, and 200 μl of the virus dilutions or infection medium was added on the cells. The cells were incubated at +37 °C for 1 h with gentle shaking every 5 min. After the incubation, 3 ml of minimum essential medium (MEM, Gibco) containing 0.5 mg/ml penicillin, 500 U/ml streptomycin, 2 mM glutamine, 2% FBS, and 1.2% Avicell was added to each well. The cells were incubated at +37 °C in a 5% CO_2_ atmosphere for 4 days, after which the medium was removed, the cells washed with PBS, and fixed with 10% formaldehyde for 30 min at room temperature. The plaques were visualised by incubating the fixed cells with 0.5% crystal violet for 10 min and washing with water. The viral titre was expressed as plaque-forming units per ml (pfu/ml).

### Virus Purification

The virus was precipitated from the cell supernatant by adding 8 % (w/v) polyethylene glycol (PEG) 8000 (Sigma-Aldrich), incubating at +4 °C with gentle mixing for 3 h, and pelleted at +4 °C, at 10,500× *g* for 50 min. The supernatant was discarded, and the pellets were carefully washed with 20 mM HEPES, 150 mM NaCl, 1 mM EDTA, pH 8.5 (HNE) and resuspended by incubating in HNE overnight at +4 °C on an orbital shaker.

The dissolved pellets were loaded onto linear 10–70 % sucrose gradients prepared in HNE. The samples were centrifuged at +4 °C at 44,500 × *g* for 17 h and the virus-containing light-scattering bands were collected. The samples were buffer exchanged to HNE using Slide-A-Lyzer dialysis cassettes (MWCO 10 kDa; ThermoFisher) at +4 °C for 1h with gentle agitation and the HNE was replaced with fresh HNE, and the dialysis continued overnight. The titre was determined using plaque titration.

### Mass spectrometry analysis of TBEV lipids

Purified virus (7×10^8^ pfu) in HNE or just HNE was mixed with methanol and chloroform (1:3:1 ratio of sample:methanol:chloroform), and vortexed for 30 s. Insoluble components were precipitated by centrifuging for 10 min at +4 °C at 20 000 × *g* and the supernatants were stored in glass vials with a nitrogen atmosphere in −80 °C.

Lipidomic profiling by UHPLC-QTOF-MS was performed at the Swedish Metabolomics Center in Umeå, Sweden. Prior to analysis the samples were transferred to microvials, evaporated under a stream of nitrogen and redissolved in 60 μl (2:1 v/v chloroform:methanol) including internal standards (tripalmitin-1,1,1-13C3, 16:0-d31 ceramide, 1,2-distearoyl-d70-sn-glycero-3-phosphocholine and 1,3-dioctadecanoyl-2-hydroxy-sn-glycerol-d5). The LC-MS analysis of the lipid extracts were performed on an Agilent 1290 Infinity UHPLC-system coupled to an Agilent 6546 Q-TOF mass spectrometer (Agilent Technologies, Waldbronn, Germany) as previously described (Diab *et al*, 2019).

The data were processed using Batch Targeted Feature Extraction algorithm within MassHunter™ ProFinder version B.10.02 (Agilent Technologies Inc., Santa Clara, CA, USA). An in-house database with exact mass and experimental retention times of lipids was used for identification.

### Statistical procedures

Statistical significance of the results were assessed either using one sample T-test (assessment of remaining F signal in 1M NaCl wash QCM-D experiments) or Welch’s T-test (all other experiments) performed in R 4.2.1.

## Supplementary figures

**Figure S1.**
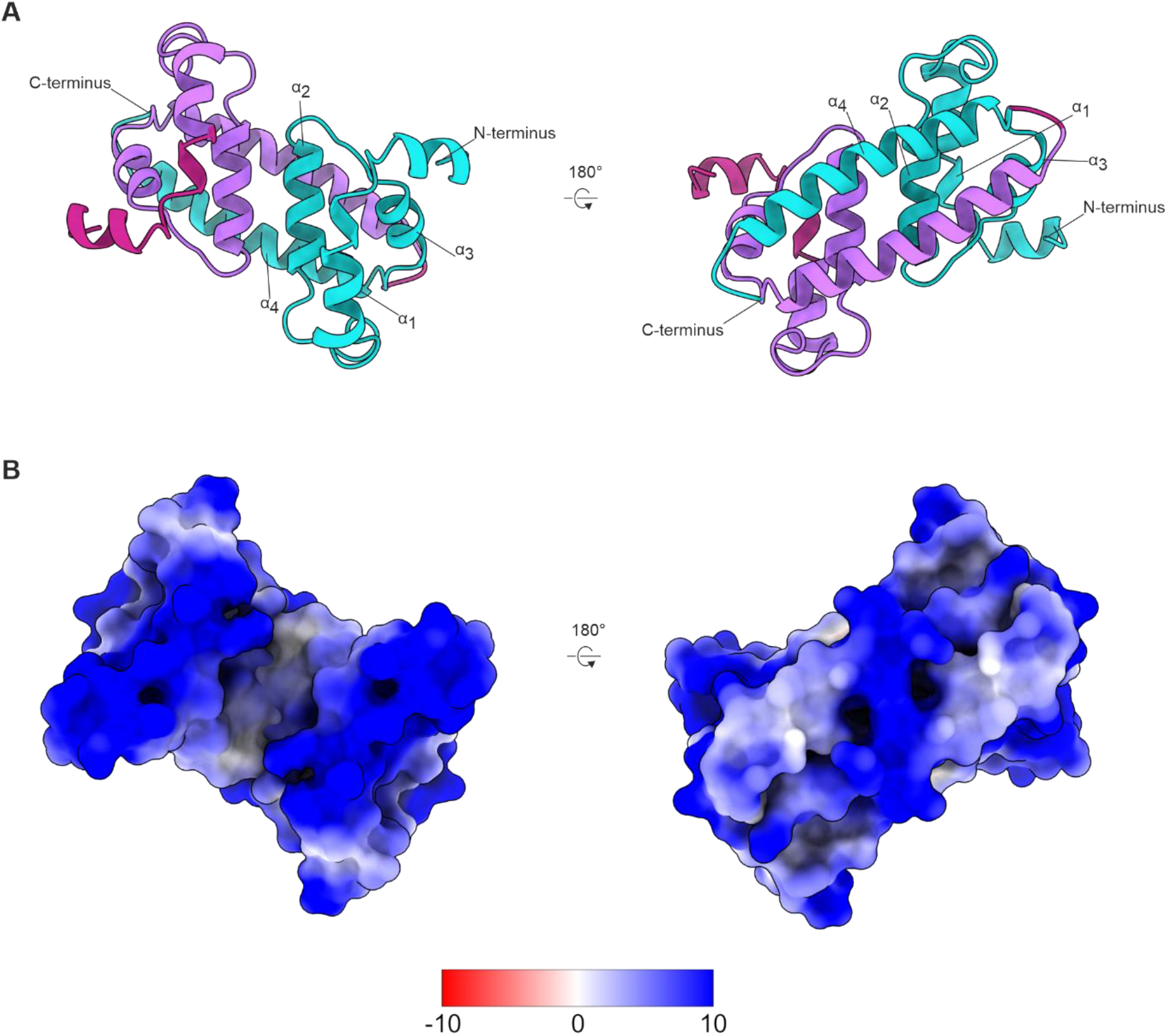
Homology model of the TBEV C protein. **A** Ribbon representation of the C homology model dimer from two angles. The chains are coloured turquoise and purple, and the α_1_–α_4_ helices and termini are labelled for one monomer. The residues truncated from the C_18–93_ construct are highlighted in magenta in the other monomer. **B** Surface representation of the C homology model dimer from the same angles as in **A** and **B**. The surfaces are coloured according to electrostatic potential according to the key (kcal / [mol × e] at 298 K). The figure was created with the homology model from Barrass *et al*. (2021) and the electrostatic potential of the surfaces were calculated in ChimeraX (Goddard *et al*, 2018)

**Figure S2.**
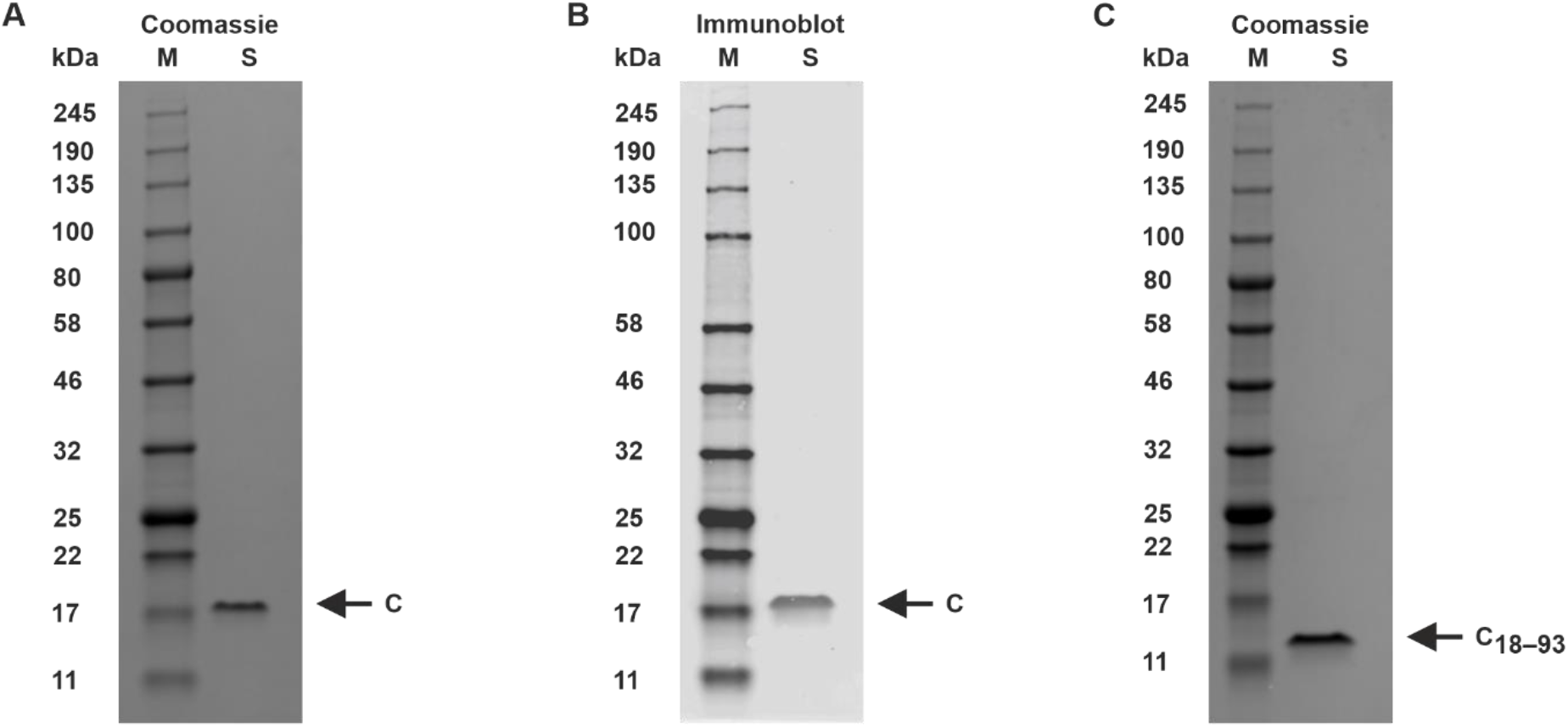
Biochemical characterisation of purified C protein **A** SDS-PAGE analysis of the purified C protein. **B** Anti-C immunoblot analysis of the purified C protein. **C** SDS-PAGE analysis of the purified C_18–93_ protein. **A-C** M lanes show the molecular size marker and S lanes the protein preparations. The sizes of the molecular size marker bands and the C, and C_18–93_ protein band positions are indicated.

**Figure S3.**
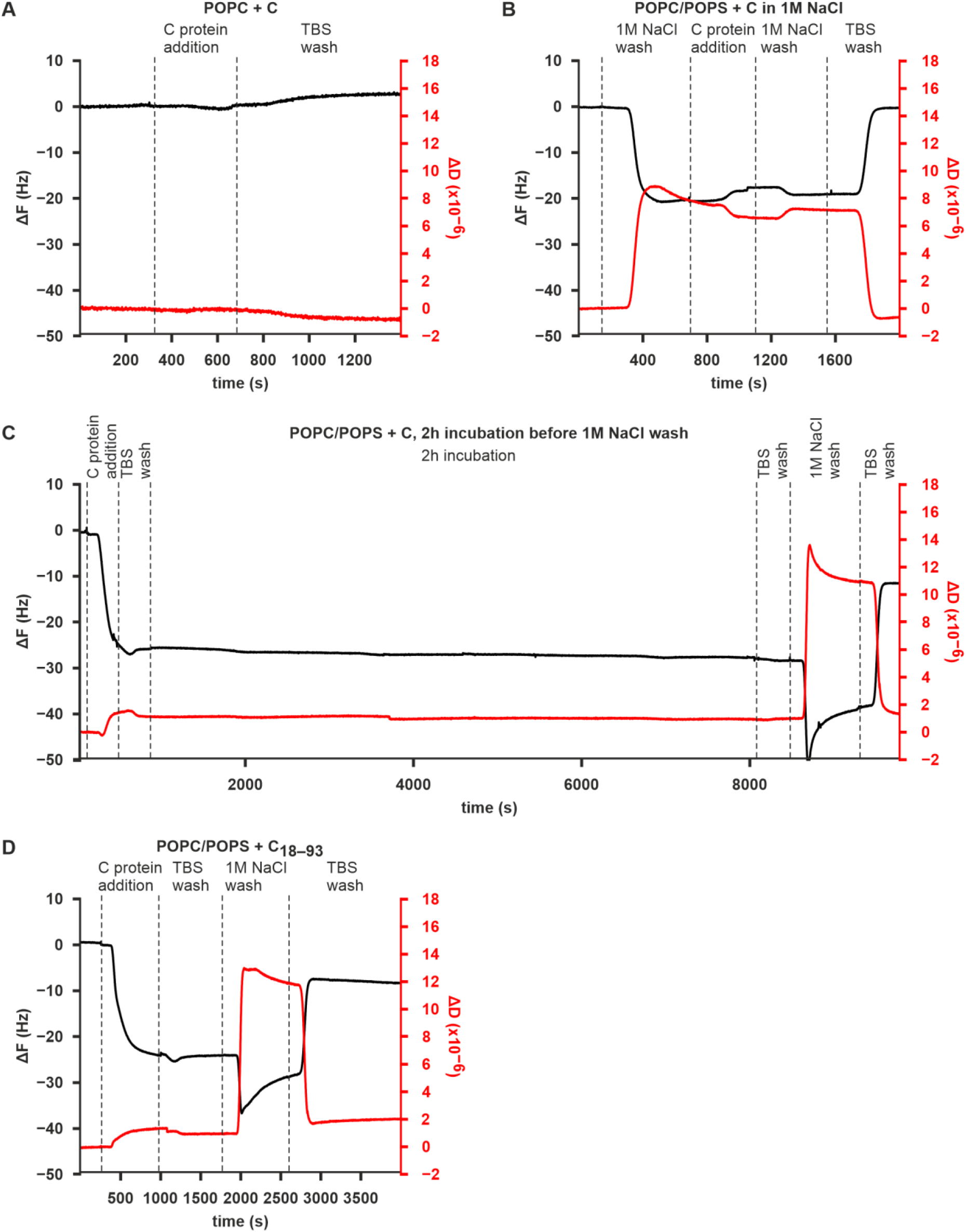
Representative QCM-D curves of the C and C_18–93_ proteins binding to SLBs. **A** Representative QCM-D curves from a C protein binding experiment on POPC SLBs. **B** Representative QCM-D curves from a C protein binding in 1M NaCl experiment on POPC/POPS SLBs. **C** Representative QCM-D curves from a C protein binding with a 1M NaCl wash after 2h incubation experiment on POPC/POPS SLBs **D** Representative QCM-D curves from a C_18–93_ protein binding experiment on POPC/POPS SLBs. Data information: In each panel, the ΔF and ΔD have been zeroed to equilibrium values after SLB formation.

**Figure S4.**
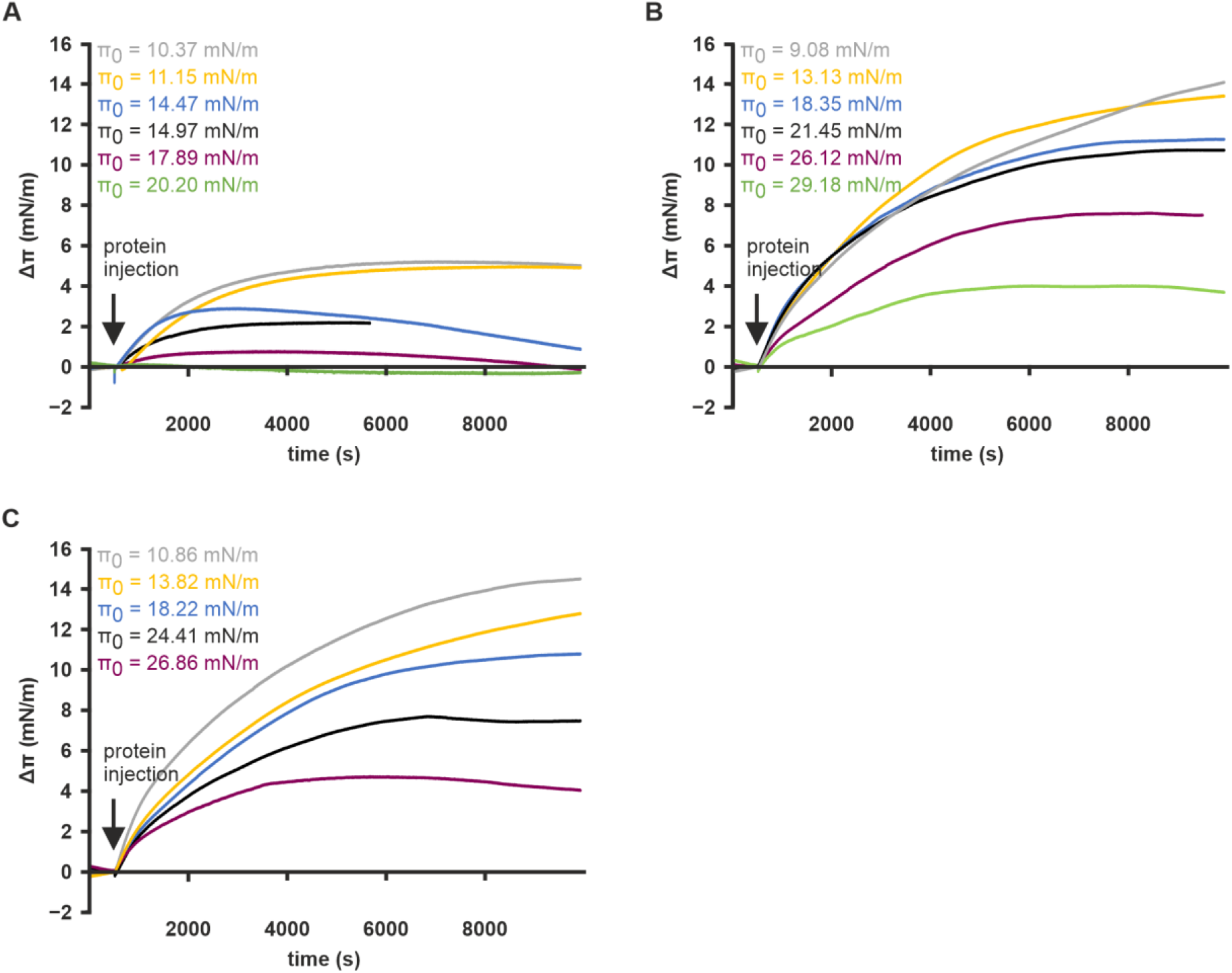
Δπ curves for C and C_18–93_ proteins injected at different Δ_0_ values shown for the same timescale as in figure 3 **A** Δπ curves for C protein injected with POPC monolayers. **B** Δπ curves for C protein injected with POPC/POPS monolayers. **C** Δπ curves for C_18–93_ protein injected with POPC/POPS monolayers. Data information: The π is zeroed to the preinjection pressure. The π_0_ values for each curve are indicated.

**Figure S5.**
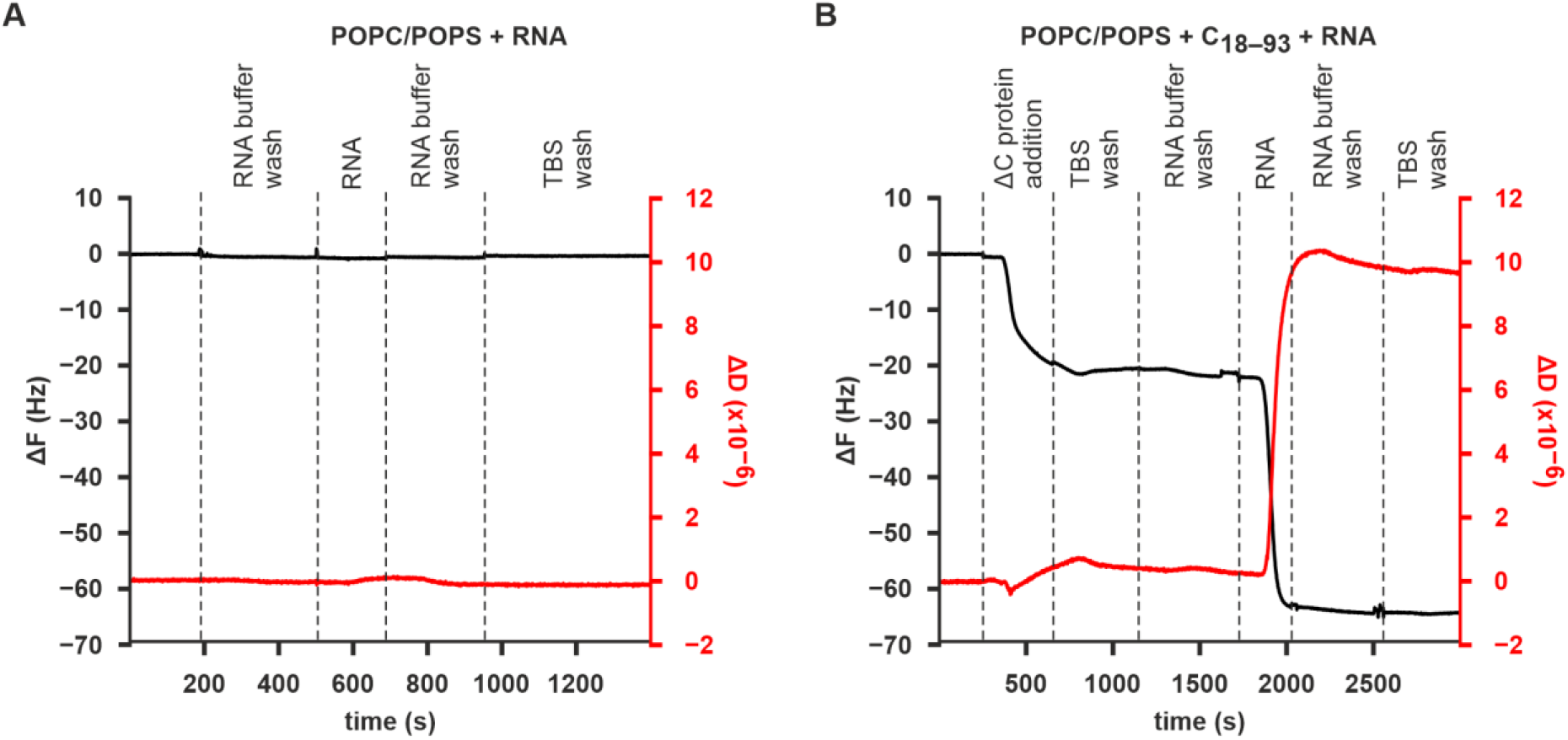
Representative QCM-D curves of RNA binding to SLBs. **A** Representative QCM-D curves from an RNA binding without protein pretreatment experiment on POPC/POPS SLBs. **B** Representative QCM-D curves from an RNA binding with pretreatment with C_18–93_ experiment on POPC/POPS SLBs. Data information: In each panel, the ΔF and ΔD have been zeroed to equilibrium values after SLB formation.

Supplementary file S1. Integrated peak areas of the lipids analysed via UHPLC-QTOF-MS

## Acknowledgements

The authors would like to thank Saana Haarma, Peter Sarin, Salla Kalaniemi, Hanna Oksanen of the Instruct-ERIC Centre Finland, Sari Korhonen, Annika Johansson, Maria Ahnlund, Ankita Shukla, Elin Larsson, Olli Vapalahti, Suvi Kuivanen, Sanna Mäki, Kang-Cheng Liu and Otto Stenberg for technical assistance, and the DNA Dream Lab facility of the Institute of Biotechnology, University of Helsinki and Konstantin Kogan personally for cloning the C protein constructs.

The research was funded by the Swedish Research Council, reference number 2018-05851 (to SJB, AKÖ, and RL); the Academy of Finland, grant number 315950 (to SJB); the Sigrid Juselius Foundation, grant number 95-7202-38 (to SJB); European Union’s Horizon 2020 Research and Innovation Programme under the Marie Skłodowska-Curie grant agreements, number 765042 (ViBRANT to SJB), and number 799929 (to MA); University of Helsinki Research Council (to LIAP); FEBS short-term fellowship (to LIAP); Swedish Research Council reference number 2021-05117 (to RL). SVB and LIAP were fellows of the Doctoral Programme in Integrative Life Science, University of Helsinki.

The Swedish Metabolomics Centre, Umeå, Sweden (www.swedishmetabolomicscentre.se) is acknowledged for the lipid profiling by UHPLC-QTOF-MS.

## Data availability

The UHPLC-QTOF-MS data have been deposited in the MetaboLights database (https://www.ebi.ac.uk/metabolights/) with the study identifier MTBLS6076 (Haug *et al*, 2020).

## Notes

https://www.ebi.ac.uk/metabolights/MTBLS6076

### Competing Interest Statement

The authors have declared no competing interest.

## References

Amberg SM, Nestorowicz A, Mccourt DW & Rice CM (1994) NS2B-3 proteinase-mediated processing in the yellow fever virus structural region: In vitro and in vivo studies. J Virol 68: 3794–3802

Ambroggio EE, Navarro GSC, Socas LBP, Bagatolli LA & Gamarnik A V. (2021) Dengue and Zika virus capsid proteins bind to membranes and self-assemble into liquid droplets with nucleic acids. J Biol Chem 297

Balinsky CA, Schmeisser H, Ganesan S, Singh K, Pierson TC & Zoon KC (2013) Nucleolin interacts with the dengue virus capsid protein and plays a role in formation of infectious virus particles. J Virol 87: 13094–13106

Barrass S V, Pulkkinen LIA, Vapalahti O, Kuivanen S, Anastasina M, Happonen L & Butcher SJ (2021) Proteome-wide cross-linking mass spectrometry to identify specific virus capsid-host interactions between tick-borne encephalitis virus and neuroblastoma cells. bioRxiv: 2021.10.29.464531

Bhuvanakantham R, Cheong YK & Ng ML (2010) West Nile virus capsid protein interaction with importin and HDM2 protein is regulated by protein kinase C-mediated phosphorylation. Microbes Infect 12: 615–625

Bhuvanakantham R & Ng ML (2013) West Nile virus and dengue virus capsid protein negates the antiviral activity of human Sec3 protein through the proteasome pathway. Cell Microbiol 15: 1688–1706

Bílý T, Palus M, Eyer L, Elsterová J, Vancová M & Růžek D (2015) Electron tomography analysis of tick-borne encephalitis virus infection in human neurons. Sci Rep 5: 1–15

Blazevic J, Rouha H, Bradt V, Heinz FX & Stiasny K (2016) The membrane anchors of the structural flavivirus proteins and their role in virus assembly. J Virol 90: JVI.00447-16

Blume A (1979) A comparative study of the phase transitions of phospholipid bilayers and monolayers. BBA - Biomembr 557: 32–44

Bogovic P & Strle F (2015) Tick-borne encephalitis: A review of epidemiology, clinical characteristics, and management. World J Clin Cases 3: 430

Boon PLS, Saw WG, Lim XX, Raghuvamsi PV, Huber RG, Marzinek JK, Holdbrook DA, Anand GS, Grüber G & Bond PJ (2018) Partial intrinsic disorder governs the dengue capsid protein conformational ensemble. ACS Chem Biol 13: 1621–1630

Calvez P, Bussières S, Éric Demers & Salesse C (2009) Parameters modulating the maximum insertion pressure of proteins and peptides in lipid monolayers. Biochimie 91: 718–733

Carvalho FA, Carneiro FA, Martins IC, Assunção-Miranda I, Faustino AF, Pereira RM, Bozza PT, Castanho MARB, Mohana-Borges R, Da Poian AT, et al (2012) Dengue virus capsid protein binding to hepatic lipid droplets (LD) is potassium ion dependent and is mediated by LD surface proteins. J Virol 86: 2096–2108

Chatel-Chaix L, Cortese M, Romero-Brey I, Bender S, Neufeldt CJ, Fischl W, Scaturro P, Schieber N, Schwab Y, Fischer B, et al (2016) Dengue virus perturbs mitochondrial morphodynamics to dampen innate immune responses. Cell Host Microbe 20: 342–356

Cheong YK & Ng ML (2011) Dephosphorylation of West Nile virus capsid protein enhances the processes of nucleocapsid assembly. Microbes Infect 13: 76–84

Cho NJ, Frank CW, Kasemo B & Höök F (2010) Quartz crystal microbalance with dissipation monitoring of supported lipid bilayers on various substrates. Nat Protoc 5: 1096–1106

Colpitts TM, Barthel S, Wang P & Fikrig E (2011) Dengue virus capsid protein binds core histones and inhibits nucleosome formation in human liver cells. PLoS One 6

Cortese M, Goellner S, Acosta EG, Neufeldt CJ, Oleksiuk O, Lampe M, Haselmann U, Funaya C, Schieber N, Ronchi P, et al (2017) Ultrastructural characterization of zika virus replication factories. Cell Rep 18: 2113–2123

Corver J, Ortiz A, Allison SL, Schalich J, Heinz FX & Wilschut J (2000) Membrane fusion activity of tick-borne encephalitis virus and recombinant subviral particles in a liposomal model system. Virology 269: 37–46

Diab J, Hansen T, Goll R, Stenlund H, Ahnlund M, Jensen E, Moritz T, Florholmen J & Forsdahl G (2019) Lipidomics in ulcerative colitis reveal alteration in mucosal lipid composition associated with the disease state. Inflamm Bowel Dis 25: 1780–1787

DiNunno NM, Goetschius DJ, Narayanan A, Majowicz SA, Moustafa I, Bator CM, Hafenstein SL & Jose J (2020) Identification of a pocket factor that is critical to Zika virus assembly. Nat Commun 11: 1–8

Dokland T, Walsh M, Mackenzie JM, Khromykh AA, Ee KH & Wang S (2004) West Nile virus core protein: Tetramer structure and ribbon formation. Structure 12: 1157–1163

Fajardo-Sánchez E, Galiano V & Villalaín J (2017) Molecular dynamics study of the membrane interaction of a membranotropic dengue virus C protein-derived peptide. J Biomol Struct Dyn 35: 1283–1294

Ferlenghi I, Clarke M, Ruttan T, Allison SL, Schalich J, Heinz FX, Harrison SC, Rey FA & Fuller SD (2001) Molecular organization of a recombinant subviral particle from tick-borne encephalitis virus. Mol Cell 7: 593–602

Fontaine KA, Leon KE, Khalid MM, Tomar S, Jimenez-Morales D, Dunlap M, Kaye JA, Shah PS, Finkbeiner S, Krogan NJ, et al (2018) The cellular NMD pathway restricts zika virus infection and is targeted by the viral capsid protein. MBio 9

Freire JM, Veiga AS, Conceição TM, Kowalczyk W, Mohana-Borges R, Andreu D, Santos NC, Da Poian AT & Castanho MARB (2013) Intracellular nucleic acid delivery by the supercharged dengue virus capsid protein. PLoS One 8

Füzik T, Formanová P, Růžek D, Yoshii K, Niedrig M & Plevka P (2018) Structure of tick-borne encephalitis virus and its neutralization by a monoclonal antibody. Nat Commun 9: 436

Gillespie LK, Hoenen A, Morgan G & Mackenzie JM (2010) The endoplasmic reticulum provides the membrane platform for biogenesis of the flavivirus replication complex. J Virol 84: 10438–10447

Goddard TD, Huang CC, Meng EC, Pettersen EF, Couch GS, Morris JH & Ferrin TE (2018) UCSF ChimeraX: Meeting modern challenges in visualization and analysis. Protein Sci 27: 14–25

Hardy JM, Newton ND, Modhiran N, Scott CAP, Venugopal H, Vet LJ, Young PR, Hall RA, Hobson-Peters J, Coulibaly F, et al (2021) A unified route for flavivirus structures uncovers essential pocket factors conserved across pathogenic viruses. Nat Commun 12

Haug K, Cochrane K, Nainala VC, Williams M, Chang J, Jayaseelan KV & O’Donovan C (2020) MetaboLights: a resource evolving in response to the needs of its scientific community. Nucleic Acids Res 48: D440–D444

Hirano M, Yoshii K, Sakai M, Hasebe R, Ichii O & Kariwa H (2014) Tick-borne flaviviruses alter membrane structure and replicate in dendrites of primary mouse neuronal cultures. J Gen Virol 95: 849–861

Katoh H, Okamoto T, Fukuhara T, Kambara H, Morita E, Mori Y, Kamitani W & Matsuura Y (2013) Japanese encephalitis virus core protein inhibits stress granule formation through an interaction with caprin-1 and facilitates viral propagation. J Virol 87: 489–502

Kaufman F, Dostálková A, Pekárek L, Thanh TD, Kapisheva M, Hadravová R, Bednárová L, Novotný R, Křížová I, Černý J, et al (2020) Characterization and in vitro assembly of tick-borne encephalitis virus C protein. FEBS Lett 594: 1989–2004

Khare B, Klose T, Fang Q, Rossmann MG & Kuhn RJ (2021) Structure of Usutu virus SAAR-1776 displays fusion loop asymmetry. Proc Natl Acad Sci U S A 118

Kiermayr S, Kofler RM, Mandl CW, Heinz FX & Messner P (2004) Isolation of capsid protein dimers from the tick-borne encephalitis flavivirus and in vitro assembly of capsid-like particles. J Virol 78: 8078–8084

Kofler RM, Heinz FX & Mandl CW (2002) Capsid protein C of tick-borne encephalitis virus tolerates large internal deletions and is a favorable target for attenuation of virulence. J Virol 76: 3534–3543

Kuivanen S, Smura T, Rantanen K, Kämppi L, Kantonen J, Kero M, Jääskeläinen A, Jääskeläinen AJ, Sane J, Myllykangas L, et al (2018) Fatal tick-borne encephalitis virus infections caused by Siberian and European subtypes, Finland, 2015. Emerg Infect Dis 24: 946–948

Kümmerer BM & Rice CM (2002) Mutations in the yellow fever virus nonstructural protein NS2A selectively block production of infectious particles. J Virol 76: 4773–4784

Larsson E, Hubert M & Lundmark R (2020) Analysis of protein and lipid interactions using liposome co-sedimentation assays. In Methods in Molecular Biology pp 119–127. Humana Press Inc.

Leung JY, Pijlman GP, Kondratieva N, Hyde J, Mackenzie JM & Khromykh AA (2008) Role of nonstructural protein NS2A in flavivirus assembly. J Virol 82: 4731–4741

Limjindaporn T, Netsawang J, Noisakran S, Thiemmeca S, Wongwiwat W, Sudsaward S, Avirutnan P, Puttikhunt C, Kasinrerk W, Sriburi R, et al (2007) Sensitization to Fas-mediated apoptosis by dengue virus capsid protein. Biochem Biophys Res Commun 362: 334–339

Lindenbach BD, Murray CL, Thiel H-J & Rice CM (2013) Flaviviridae. In Fields Virology, Knipe D & Howley P (eds) Philadelphia: Lippincott Williams & Wilkins

Liu K-C, Pace H, Larsson E, Hossain S, Kabedev A, Shukla A, Jerschabek V, Mohan J, Bergström CAS, Bally M, et al (2022) Membrane insertion mechanism of the caveola coat protein Cavin1. Proc Natl Acad Sci 119

Liu WJ, Chen HB & Khromykh AA (2003) Molecular and functional analyses of Kunjin virus infectious cDNA clones demonstrate the essential roles for NS2A in virus assembly and for a nonconservative residue in NS3 in RNA replication. J Virol 77: 7804–7813

Lorenz IC, Kartenbeck J, Mezzacasa A, Allison SL, Heinz FX & Helenius A (2003) Intracellular assembly and secretion of recombinant subviral particles from tick-borne encephalitis virus. J Virol 77: 4370–4382

Ma L, Jones CT, Groesch TD, Kuhn RJ & Post CB (2004) Solution structure of dengue virus capsid protein reveals another fold. Proc Natl Acad Sci U S A 101: 3414–3419

Ma L, Li F, Zhang J-W, Li W, Zhao D-M, Wang H, Hua R-H & Bu Z-G (2018) Host factor SPCS1 regulates the replication of Japanese encephalitis virus through interactions with transmembrane domains of NS2B. J Virol: JVI.00197-18

Markoff L, Falgout B & Chang A (1997) A Conserved internal hydrophobic domain mediates the stable membrane integration of the dengue virus capsid protein. Virology 233: 105–117

Marsh D (1996) Lateral pressure in membranes. Biochim Biophys Acta - Rev Biomembr 1286: 183–223

Martín-Acebes MA, Merino-Ramos T, Blázquez A-B, Casas J, Escribano-Romero E, Sobrino F & Saiz J-C (2014) The composition of West Nile virus lipid envelope unveils a role of sphingolipid metabolism in flavivirus biogenesis. J Virol 88: 12041–12054

McMahon HT & Boucrot E (2015) Membrane curvature at a glance. J Cell Sci 128: 1065–1070

Mebus-Antunes NC, Ferreira WS, Barbosa GM, Neves-Martins TC, Weissmuller G, Almeida FCL & Da Poian AT (2022) The interaction of dengue virus capsid protein with negatively charged interfaces drives the in vitro assembly of nucleocapsid-like particles. PLoS One 17: e0264643

Van Meer G, Voelker DR & Feigenson GW (2008) Membrane lipids: Where they are and how they behave. Nat Rev Mol Cell Biol 9: 112–124

Miller S, Kastner S, Krijnse-Locker J, Bühler S & Bartenschlager R (2007) The non-structural protein 4A of dengue virus is an integral membrane protein inducing membrane alterations in a 2K-regulated manner. J Biol Chem 282: 8873–8882

Miorin L, Romero-Brey I, Maiuri P, Hoppe S, Krijnse-Locker J, Bartenschlager R & Marcello A (2013) Three-dimensional architecture of tick-borne encephalitis virus replication sites and trafficking of the replicated RNA. J Virol 87: 6469–6481

Nemésio H, Palomares-Jerez F & Villalaín J (2011) The membrane-active regions of the dengue virus proteins C and E. Biochim Biophys Acta - Biomembr 1808: 2390–2402

Nemésio H, Palomares-Jerez MF & Villalaín J (2013) Hydrophobic segment of dengue virus C protein. Interaction with model membranes. Mol Membr Biol. 30: 273–287

Offerdahl DK, Dorward DW, Hansen BT & Bloom ME (2012) A three-dimensional comparison of tick-borne flavivirus infection in mammalian and tick cell lines. PLoS One 7

Panayiotou C, Lindqvist R, Kurhade C, Vonderstein K, Pasto J, Edlund K, Upadhyay AS & Överby AK (2018) Viperin restricts Zika virus and tick-borne encephalitis virus replication by targeting NS3 for proteasomal degradation. J Virol: JVI.02054-17

Patkar CG & Kuhn RJ (2008) Yellow fever virus NS3 plays an essential role in virus assembly independent of its known enzymatic functions. J Virol 82: 3342–3352

Peña J & Harris E (2012) Early dengue virus protein synthesis induces extensive rearrangement of the endoplasmic reticulum independent of the UPR and SREBP-2 pathway. PLoS One 7: 1–15

Perera R, Riley C, Isaac G, Hopf-Jannasch AS, Moore RJ, Weitz KW, Pasa-Tolic L, Metz TO, Adamec J & Kuhn RJ (2012) Dengue virus infection perturbs lipid homeostasis in infected mosquito cells. PLoS Pathog 8

Poonsiri T, Wright GSA, Solomon T & Antonyuk S V. (2019) Crystal structure of the Japanese encephalitis virus capsid protein. Viruses 11

Pulkkinen LIA, Barrass S V, Domanska A, Överby AK, Anastasina M & Butcher SJ (2022) Molecular organisation of tick-borne encephalitis virus. Viruses 14: 1–17

Pulkkinen LIA, Butcher S & Anastasina M (2018) Tick-borne encephalitis virus: A structural view. Viruses 10: 350

Reid CR & Hobman TC (2017) The nucleolar helicase DDX56 redistributes to West Nile virus assembly sites. Virology 500: 169–177

Renner M, Dejnirattisai W, Carrique L, Martin IS, Karia D, Ilca SL, Ho SF, Kotecha A, Keown JR, Mongkolsapaya J, et al (2021) Flavivirus maturation leads to the formation of an occupied lipid pocket in the surface glycoproteins. Nat Commun 12: 1–9

Ruzek D, Avšič Županc T, Borde J, Chrdle A, Eyer L, Karganova G, Kholodilov I, Knap N, Kozlovskaya L, Matveev A, et al (2019) Tick-borne encephalitis in Europe and Russia: Review of pathogenesis, clinical features, therapy, and vaccines. Antiviral Res 164: 23–51

Samsa MM, Mondotte JA, Iglesias NG, Assunção-Miranda I, Barbosa-Lima G, Da Poian AT, Bozza PT & Gamarnik A V. (2009) Dengue virus capsid protein usurps lipid droplets for viral particle formation. PLoS Pathog 5

Samuel GH, Wiley MR, Badawi A, Adelman ZN & Myles KM (2016) Yellow fever virus capsid protein is a potent suppressor of RNA silencing that binds double-stranded RNA. Proc Natl Acad Sci 113: 13863–13868

Schindelin J, Arganda-Carreras I, Frise E, Kaynig V, Longair M, Pietzsch T, Preibisch S, Rueden C, Saalfeld S, Schmid B, et al (2012) Fiji: An open-source platform for biological-image analysis. Nat Methods 9: 676–682

Shang Z, Song H, Shi Y, Qi J & Gao GF (2018) Crystal structure of the capsid protein from Zika virus. J Mol Biol 430: 948–962

Simmonds P, Becher P, Bukh J, Gould EA, Meyers G, Monath T, Muerhoff S, Pletnev A, Rico-Hesse R, Smith DB, et al (2017) ICTV virus taxonomy profile: Flaviviridae. J Gen Virol 98: 2–3

Slomnicki LP, Chung DH, Parker A, Hermann T, Boyd NL & Hetman M (2017) Ribosomal stress and Tp53-mediated neuronal apoptosis in response to capsid protein of the Zika virus. Sci Rep 7

Stadler K, Allison SL, Schalich J & Heinz FX (1997) Proteolytic activation of tick-borne encephalitis virus by furin. J Virol 71: 8475–8481

Tabata K, Arimoto M, Arakawa M, Suzuki R, Matsuura Y & Morita E (2016) Unique requirement for ESCRT factors in flavivirus particle formation on the endoplasmic reticulum. Cell Rep 16

Tan TY, Fibriansah G, Kostyuchenko VA, Ng TS, Lim XX, Zhang S, Lim XN, Wang J, Shi J, Morais MC, et al (2020) Capsid protein structure in Zika virus reveals the flavivirus assembly process. Nat Commun 11

Thorsen MK, Lai A, Lee MW, Hoogerheide DP, Wong GCL, Freed JH & Heldwein EE (2021) Highly basic clusters in the herpes simplex virus 1 nuclear egress complex drive membrane budding by inducing lipid ordering. MBio 12

Urbanowski MD & Hobman TC (2013) The West Nile virus capsid protein blocks apoptosis through a phosphatidylinositol 3-kinase-dependent mechanism. J Virol 87: 872–881

Vonderstein K, Nilsson E, Hubel P, Nygård Skalman L, Upadhyay A, Pasto J, Pichlmair A, Lundmark R & Överby AK (2018) Viperin targets flavivirus virulence by inducing assembly of noninfectious capsid particles. J Virol 92

Voßmann S, Wieseler J, Kerber R & Kümmerer BM (2015) A basic cluster in the n terminus of yellow fever virus ns2a contributes to infectious particle production. J Virol 89: 4951–4965

Wang HY, Bharti D & Levental I (2020) Membrane heterogeneity beyond the plasma membrane. Front Cell Dev Biol 8: 580814

Warburg O & Christian W (1942) Isolation and crystallization of enolase. Biochem Z 310: 384–421

Xie X, Gayen S, Kang C, Yuan Z & Shi P-Y (2013) Membrane topology and function of dengue virus NS2A protein. J Virol 87: 4609–4622

Xie X, Zou J, Puttikhunt C, Yuan Z & Shi P-Y (2015) Two distinct sets of NS2A molecules are responsible for dengue virus RNA synthesis and virion assembly. J Virol 89: 1298–1313

Xu Z, Anderson R & Hobman TC (2011) The capsid-binding nucleolar helicase DDX56 is important for infectivity of West Nile virus. J Virol 85: 5571–5580

Yang JS, Ramanathan MP, Muthumani K, Choo AY, Jin SH, Yu QC, Hwang DS, Choo DK, Lee MD, Dang K, et al (2002) Induction of inflammation by West Nile Virus capsid through the caspase-9 apoptotic pathway. Emerg Infect Dis 8: 1379–1384

Yau W-L, Nguyen-Dinh V, Larsson E, Lindqvist R, Överby AK & Lundmark R (2019) Model system for the formation of tick-borne encephalitis virus replication compartments without viral RNA replication. J Virol 93

Yu C, Achazi K, Möller L, Schulzke JD, Niedrig M & Bücker R (2014) Tick-borne encephalitis virus replication, intracellular trafficking, and pathogenicity in human intestinal Caco-2 cell monolayers. PLoS One 9: 1–10

